# Identification of novel origins of transfer across bacterial plasmids

**DOI:** 10.1101/2024.01.30.577996

**Authors:** Manuel Ares-Arroyo, Amandine Nucci, Eduardo P.C. Rocha

## Abstract

Conjugative plasmids are important drivers of bacterial evolution, but most plasmids lack genes for conjugation. It is currently not known if the latter can transfer because origins of transfer by conjugation (*oriT*), which would allow their mobilization by conjugative plasmids, are poorly known. Here, we identify and characterize occurrences of known *oriT* families across thousands of plasmids confirming that most conjugative and mobilizable plasmids still lack identifiable families of *oriTs*. They reveal clear patterns in terms of intergenic position, distance to the relaxases, and MOB-type association. This allowed to develop a computational method to discover novel *oriT*s. As a proof of concept, we identify 21 novel *oriTs* from the nosocomial pathogens *Escherichia coli*, *Klebsiella pneumoniae*, and *Acinetobacter baumannii*, some of them responsible for the mobility of critical antimicrobial resistance genes. These 21 *oriT* families share key characteristics of the others and fill most of the missing diversity of *oriTs* in relaxase-encoding plasmids both in terms of frequency and phylogeny. We confirmed experimentally the function of six of them. The ability to identify novel *oriT*s paves the way to explore conjugation across bacterial plasmids, notably among the majority lacking conjugation-related genes.

## INTRODUCTION

Horizontal gene transfer has a key role in providing novel functions to bacteria^1^. Conjugation is one of its major mechanisms^2^, driving the spread of antibiotic resistance, rhizobial traits, or complex virulence factors^3^. Conjugation is driven by mobile genetic elements that integrate the chromosome (ICEs) or remain extra-chromosomal (plasmids)^4^. It requires three components^5^: an origin of transfer (*oriT*), which is a DNA sequence recognized by the relaxase (MOB), which produces a single strand cut at the *nic* site of the *oriT* and attaches to it. The nucleoprotein complex then interacts with the mating pair formation (MPF) apparatus that transfers it directly into another cell. Once in the recipient cell, the ssDNA is ligated and replicated, thereby effectively replicating the MGE remaining in the donor cell. The *oriT*, MOB and MPF often co-localize in plasmids and ICEs^6–8^, but exceptions have been described^9–11^.

Early results showed that only about a quarter of all plasmids are conjugative, *i.e.* encode all protein coding genes required for autonomous conjugation (MPF and MOB)^5^. Another quarter encode just a relaxase and use an MPF produced by a conjugative element. This leaves around half of the plasmids as putatively non-transmissible, which is at odds with the idea that plasmid transmission is important for their long-term survival^12,13^. The hypothesis of non-transmissibility of so many plasmids is also at odds with the observation that most of them, conjugative or not, have very scattered distributions within species, suggesting they are frequently transferred^14,15^. While this matter remained a mystery for close to a decade, recent work suggests that many putatively non-transmissible plasmids can be transferred^16^. About 7% of the plasmids are phages (phage-plasmids) and transfer within viral particles^17^. Rolling-circle replication plasmids might not require relaxases nor *oriTs*, since their replication protein could act as a coupling protein, providing a single-stranded copy of the plasmid to the MPF apparatus^18^. Finally, plasmids with an *oriT* using relaxases and MPFs encoded in one or two other MGEs^19–21^ might be more frequent than previously thought, e.g. they are ∼50% of *S. aureus* plasmids^22,23^.

The recent advances in understanding plasmid mobility in model species raise the question of the mobility of plasmids by conjugation in the others. Unfortunately, the known *oriT* families are restricted to only a few species, and it is not known how frequently they can be found in other clades. The lack of recognizable *oriTs* hampers the understanding of the patterns of transfer of plasmids and integrative mobilizable elements. Breaking this deadlock would allow to identify the transmissible plasmids and to unravel their functional dependencies to the other plasmids. It would also provide novel vector tools for the molecular manipulation of non-model species.

Here, we start by searching for sequences similar to known *oriT*s in bacterial plasmids. Among plasmids lacking *oriT*, many encode conjugative systems or relaxases and should therefore have an *oriT*. We characterize these occurrences in terms of genetic context (proximity to relaxase, intergenic region, relaxase and plasmid type) to understand where to look for novel *oriTs*. Using this information, we were able to identify novel families of *oriTs* in many of the conjugative and mobilizable plasmids that lacked them. As a proof of concept, we identified 21 novel families of *oriTs* in three model species (*E. coli*, *K. pneumoniae*, *A. baumannii*). We used computational approaches to test the pertinence of the novel *oriTs* in terms of properties that were not used to identify them. Finally, we made an experimental proof of concept that some of the novel *oriT* families allow plasmid transfer in *E. coli* using an heterologous system of conjugation^24^. This shows that our method has the potential to unravel many of the unknown origins of transfer by conjugation.

## RESULTS

### Most plasmids lack a recognizable origin of transfer

It was previously shown that most plasmids in *E. coli* and *S. aureus* have *oriTs*^22,23^. Here, we searched for all known types of *oriTs* across all Bacteria (38,057 plasmids from 32,798 complete genomes). We found 11,410 *oriTs* in 9,683 plasmids (25%) (Figure S1A). Very few plasmids have more than one *oriT* from a given family (Figure S1B and S1C). We then classed the plasmids into those: encoding the key genes necessary for conjugation (conjugative plasmids: pCONJ), encoding a few MPF genes (decayed conjugative plasmids: pdCONJ), encoding a relaxase (pMOB), and lacking a relaxase (pMOBless) (Figure 1A). The known *oriTs* were found in less than 45% of the pCONJ and are even rarer among pdCONJ (35%) and pMOB (28%) (Figure 1B-D). The lack of *oriTs* is even more striking in the pMOBless, once phage-plasmids are removed^17^, since only 16% have an *oriT* (Figure 1E). Hence, the currently known *oriT* families only match 36% of the plasmids that should have one and miss most of the others.

**Figure 1.**
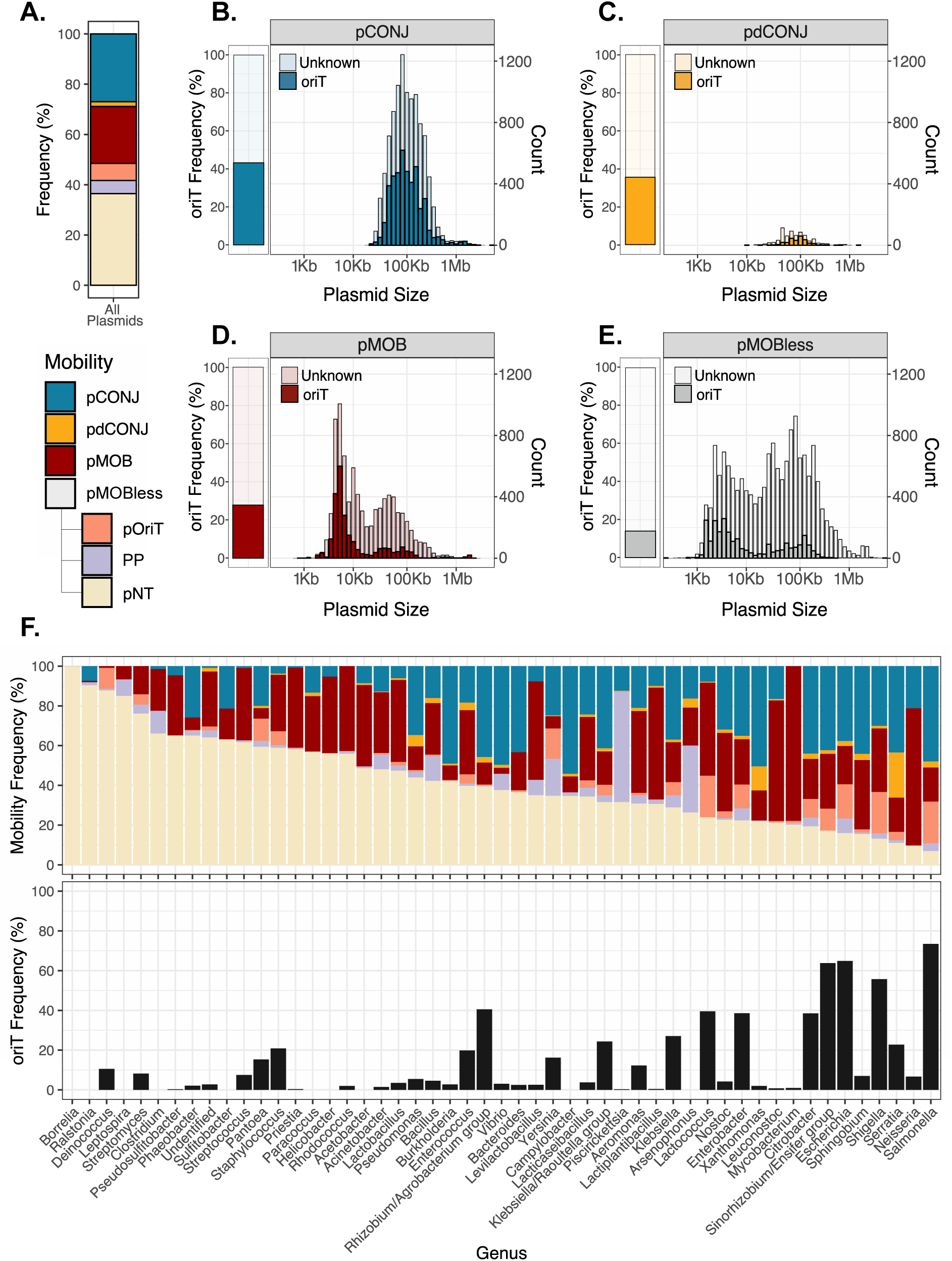
**A.** Plasmid distribution according to their mobility. **B-E.** Per conjugative type (pCONJ, pdCONJ, pMOB and pMOBless), fraction of plasmids in which an *oriT* has been identified (left), and distribution of plasmid size (right). The phage-plasmids were excluded from the set of pMOBless. **F.** Conjugative characteristics of the genera with more than 100 plasmids in RefSeq. The upper figure represents the frequency of each mobility type (pCONJ, pdCONJ, pMOB, pOriT, PP and pNT). The bottom figure represents the frequency of all the plasmids in which a known *oriT* has been identified. The legend is at the mid-left of the figure.

We then characterized these plasmids. The distribution of sizes of plasmids reveal few surprises^25^, with pCONJ and pdCONJ being larger than the others (Figure 1B). Size distributions are similar whether the plasmids have a recognizable *oriT* or not, suggesting that the two groups of plasmids are not fundamentally different (and that we are missing the *oriTs*). The pMOB and pMOBless lacking *oriT* tend to be larger than those having one, possibly because of the presence of large plasmids from clades whose conjugation machineries remain yet to be characterized (e.g. Phyla Actinomycetota, Deinococcota, or Spirochaetota) (Figure 1F, Figure S2). Yet, many oriTs are missed in clades where the conjugation machineries are identifiable. In genus like *Neisseria*, *Sphingobium* or *Leuconostoc* more than 80% of the plasmids could be assigned a mechanism of mobilization by conjugation (pCONJ or pMOB), but we lack identifiable *oriTs* in up to 95% of their plasmids. These results show that we ignore most *oriTs*.

### Known origins of transfer share key characteristics

Since we miss most *oriT*s, we studied the characteristics of the known ones to subsequently devise an approach to discover novel families. The vast majority of *oriTs* (93%) are in intergenic regions, even if these are less than 20% of the plasmid sequences (Figure 2A, Figure S3). Intergenic regions with *oriT*s are larger (median (M) = 496 bp) than the others (M = 206 bp) (p<0.001, Wilcoxon test, Figure 2B). This could be explained by the presence of the *oriT*-associated *nic* site plus the protein-binding domains for the relaxase and relaxosome-accessory proteins in these regions. Accordingly, we observe some differences according to the MOB type associated to the *oriT* (Figure 2C). Of note, these intergenic regions sometimes accommodate more than the *oriT*, e.g. in ColE1-like plasmids they may include the non-coding RNAs in charge of replication initiation^26^.

**Figure 2.**
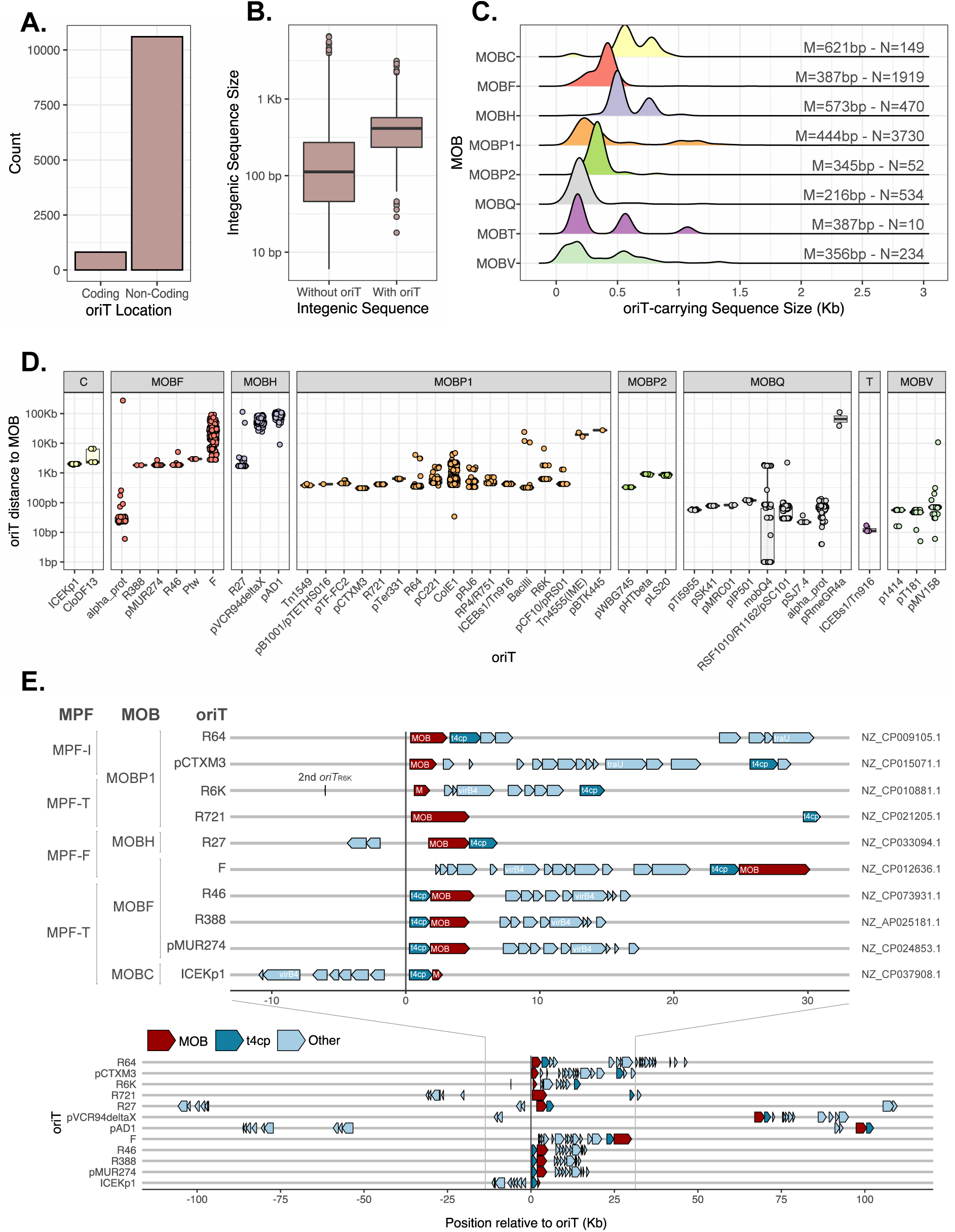
**A.** Location of the *oriT* regarding coding and non-coding sequences. **B.** Size of the intergenic sequences according to the presence/absence of *oriTs*. All sequences belong to plasmids with known *oriTs*. **C.** Size of the intergenic sequences carrying an *oriT* depending on their MOB type. M=Mean. N=Sample size. **D.** Distance between each *oriT* family and MOB type. **E.** Location of conjugative genes identified with CONJscan relative to the *oriT* identified in pCONJ from *E. coli*. The lower plot of the figure represents the position of every conjugative gene, whereas the upper plot zooms into the closer positions to the *oriT* (−10 Kb to +30 Kb, being the *oriT* in position 0). The name of the *oriT*, MOB and MPF type are indicated at the left of the figure. The accession number of the plasmid represented is at the right. Arrows represent conjugative genes, being the red one the MOB, the dark blue the coupling protein, and the light blue the remaining genes. To note, examples of *oriT*_pAD1-H_ and *oriT*_pVCR94X_ are only represented in the bottom part since the MOB is far from the *oriT*.

In MGEs, one typically finds DNA motifs and their interacting proteins in strong genetic linkage. This explains why many *oriTs* were firstly described close to the relaxase gene^6–8^. However, some *oriTs* were found kilobases away from the latter^9,10^. A systematic characterization of the location of the known *oriTs* in relation to the cognate relaxases revealed that one family of *oriT* is usually associated with one single MOB type (canonical *oriT*-MOB pair). The few (3%, Figure S4) exceptions may be due to recent changes in the plasmid’s gene repertoires and were excluded from further analysis. The distance between the *oriT* and the relaxase is usually very small (median = 628 bp), being the *oriT* upstream to the relaxase in most cases (Figure 2D and S5). The variations are due to the specific and conserved location of the relaxases within the relaxase operon for a given MOB type (Figure 2E). For example, MOB_Q_, MOB_T_ or MOB_V_ are usually just adjacent to the *oriT*. In contrast, MOB_C_ and MOB_F_ are usually preceded by the coupling protein, therefore located at ∼2Kb and ∼1.5Kb, respectively (discarding the exceptional *oriT*_F_). Distances between the *oriT* and the relaxases were large in only three well-represented cases: *oriT*_F_ (∼30Kb opposite to the MOB_F_, at opposite edges of the MPF locus), *oriT*_pVCR941′X_ and *oriT*_pAD1-H_ (∼50Kb apart from the MOB_H_). Interestingly, all of these are associated to MPF type F (Figure 2E). Altogether, these analyses show that the location of *oriTs* obeys to certain rules that may be used to search for novel ones.

### Identification of novel oriTs in E. coli, K. pneumoniae, and A. baummanii

Based on our results, *oriTs* are expected to be in large non-coding regions close to the relaxase. We therefore picked the closest intergenic sequence upstream to the relaxase that was large enough to harbor an *oriT* in the plasmids of three species with many multidrug resistant strains from the ESKAPEE pathogens group. *Escherichia coli* is a species in which we know most of the *oriTs* (∼80% of 3,689 pCONJ/pMOB). *Klebsiella pneumoniae* is from a closely related genus but we ignore most of its *oriTs* (∼35% of 3,006 pCONJ/pMOB have an *oriT*). In *Acinetobacter baumannii* we ignore almost all *oriTs* (∼1% of 336 pCONJ/pMOB). We retrieved 1,133 non-redundant non-coding sequences preceding the relaxase of the plasmids of these species (710, 332, and 91, respectively) and clustered them according to their sequence similarity (Figure 3A). The presence of known *oriTs* in some of these plasmids allows to compare their clusters with those produced for putative novel *oriT*s. Among the 1,133 sequences, 809 were split into 41 clusters and 324 were discarded (small clusters or singletons). The examination of the 41 clusters revealed 21 (220 sequences) matching plasmids lacking known *oriT*s (Figure 3B-C). These are the most promising candidate novel *oriT* regions (File S1).

**Figure 3.**
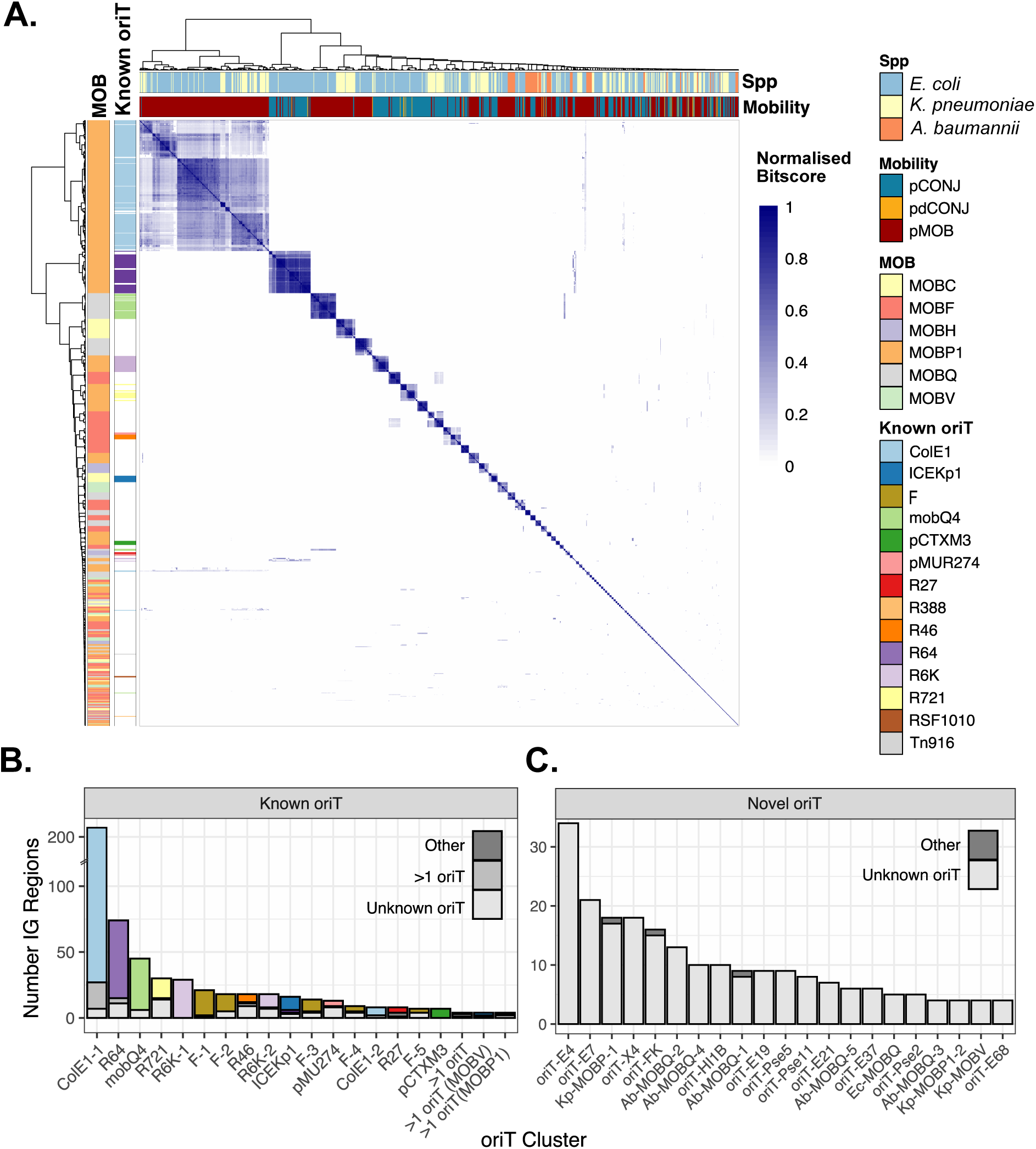
**A.** Clustering analysis of the intergenic sequences upstream to the MOB from the three species. Colors at the top of the heatmap represent the species and mobility of the plasmid, whereas colors at the left of the heatmap represent the already known *oriT* and MOB associated to the non-coding sequence. The legend is at the right of the figure. **B.** and **C.** Clusters inferred from the analysis of intergenic (IG) regions. The 20 clusters with known *oriTs* are represented at the left (B.) and the novel *oriTs* at the right (C.), being the colors indicating the *oriT* (same legend as 3A).

The clusters of novel and known *oriTs* show similar characteristics (Figure S6). First, they are specifically associated to groups of pCONJ or pMOB (but not both). Secondly, the plasmids of a given *oriT* family tend to have a distribution of size more homogeneous than the rest of the plasmids (Figure S6B). Finally, their sequences are located at the expected position relative to the relaxases for each MOB type. Interestingly, sequences of *E. coli* and *K. pneumoniae* are intermingled within a few clusters (e.g. *oriT*_ColE1_ and *oriT*_ICEKp1_), suggesting the existence of families of plasmids with the same *oriT* in both species^27^. In contrast, the *oriTs* of the more distantly related *A. baumannii* were almost always species-specific (Figure 3A, Figure S6C).

This host specificity explains the difficulties encountered when looking for *oriTs* via sequence similarity when one ignores all the *oriTs* of almost all bacterial species. They can only be discovered by *ab initio* methods.

Our method to identify origins of transfer does not search for key characteristics of *oriTs* that we can now use to test its reliability. (I) The *nic* site is the short (<10 nt) sequence located within the *oriT* that is recognized and cleaved by the relaxase. We screened for known *nic* sites in the clusters with novel *oriTs*, finding similar sequences in 8 of the putative new *oriT* families. They were all associated to the MOB types of the original *nic* sites (Figure S7). (II) *OriTs* tend to exhibit DNA secondary structures close to the *nic* site^28^. Indeed, there were various putative DNA hairpins in almost all the sequences, especially next to the putative *nic* sites (Figure S7). (III) Finally, pairs of plasmids with similar relaxases (>50% identity) tend to have the same type of *oriT* (Figure 4A). This is also the case for pairs of plasmids with the same type of novel *oriT*, whose relaxases are usually more than 60% identical (Figure 4A).

**Figure 4.**
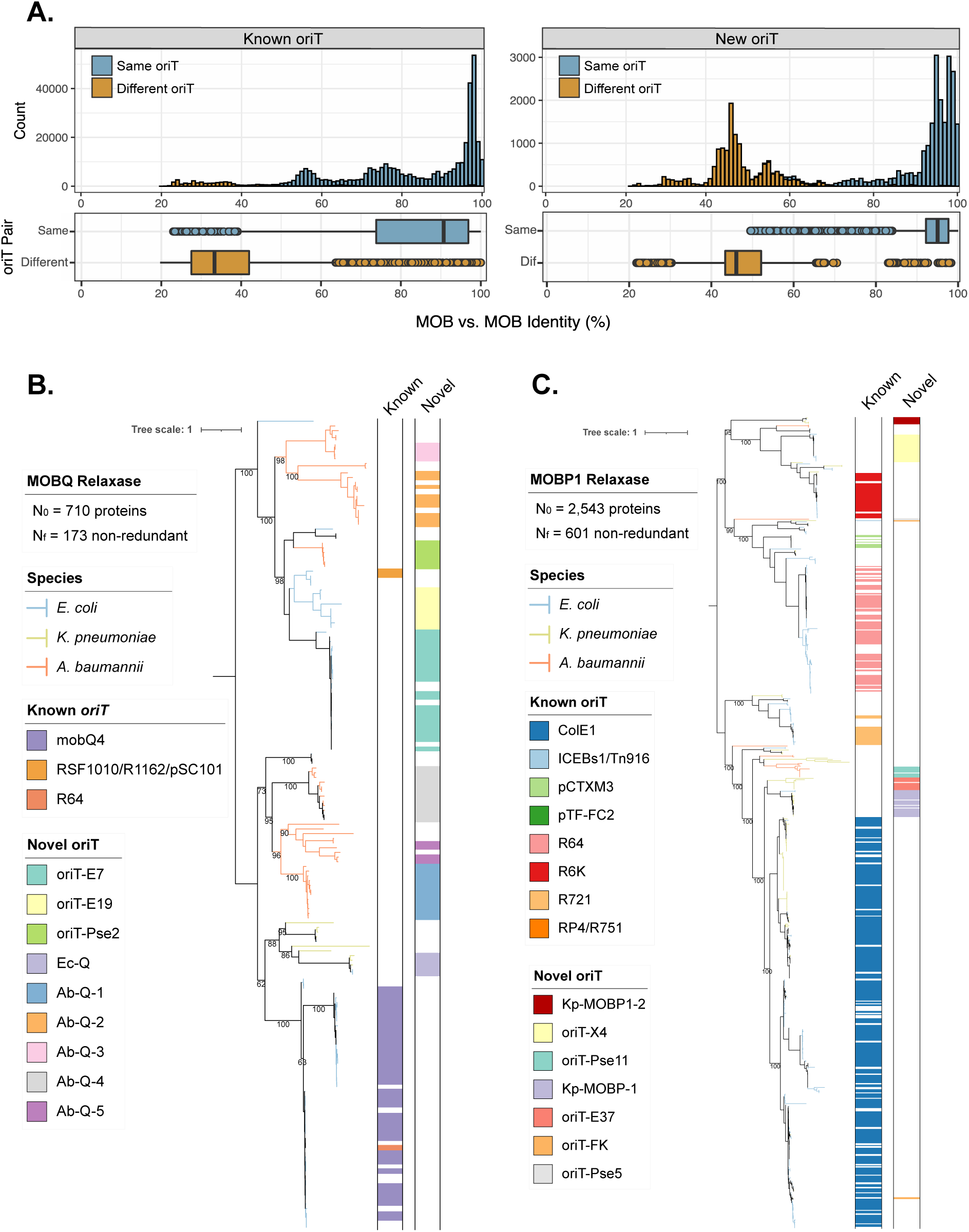
**A.** Distribution of the similarity between relaxases when they are associated to the same or to different *oriT*. The left graph shows the values for previously known *oriT*s and the one on the right to the novel ones. Relaxases with unknown *oriTs* have been discarded from the plots. **B.** and **C.** Phylogenetic trees of the MOB_Q_ (B) and MOB_P1_ (C). The colors of the branches indicate the species (*E. coli*, *K. pneumoniae* and *A. baumannii*). The color bars at the right of the trees indicate the known *oriT* (left) and novel *oriT* family (right) associated to the relaxase. The legend is at the left of each tree. Bootstraps values close to the tips of the tree have been removed for better visualization. They are available in File S2 and S3.

The phylogenies of the relaxases show that the novel putative *oriTs* complement the distribution of the previously known ones, *i.e.*, they are associated to specific branches of the tree, covering almost all the MOB diversity (Figure 4B and 4C). In the clades of relaxases where there were known *oriTs* we do not identify novel ones. In the others, we can now fill the knowledge gap and identify putative *oriT*s. The few plasmids still lacking *oriT*s are scattered across the tree. They are often in clades where either all plasmids are nearly identical or are very different and our method cannot yet reliably identify *oriT*s in such cases. Together, these analyses suggest that the 21 clusters represent new families of putative *oriTs* in the three species.

### Newly identified *oriTs* are functional in wild-type populations

To test our method, 9 novel *oriT*s responsible for the spread of AMR were synthesized and inserted into the non-transmissible cloning vector pCOLADuet^TM^-1 (Figure 5A, Table S1). In the wild-type plasmids associated to the MOB_Q_, we identified one/two genes upstream the novel *oriTs* in the opposite direction to the MOB that could be accessory genes of the relaxosome since their homologs were identified as such in some IncQ/MOB_Q_ plasmids^29,30^. Thus, a third construct including them was built for MOB_Q_ plasmids (Figure 5A and S8). Each vector was independently transformed into *E. coli* MFD*pir*, which has a chromosomal-encoded RP4 conjugative machinery. Hence, our experiment will only work if the *oriT*s and relaxases can be mobilized by this specific conjugative system. Because we were expecting low conjugation rates in this artificial setup, which serves as a proof of principle, we made long mating experiments (24h) into *E. coli* 10B. We recovered transconjugants for 6 out of the 9 *oriTs* (confirmed by PCR, Figure 5B), including two originating from *E. coli* plasmids (*oriT*_E7_ and *oriT*_X4_), three from *A. baumannii*, and one from *K. pneumoniae*. No transconjugants were observed for the negative controls (original pCOLADuet^TM^-1 and plasmid with relaxases but no *oriT,* Figure S8).

**Figure 5.**
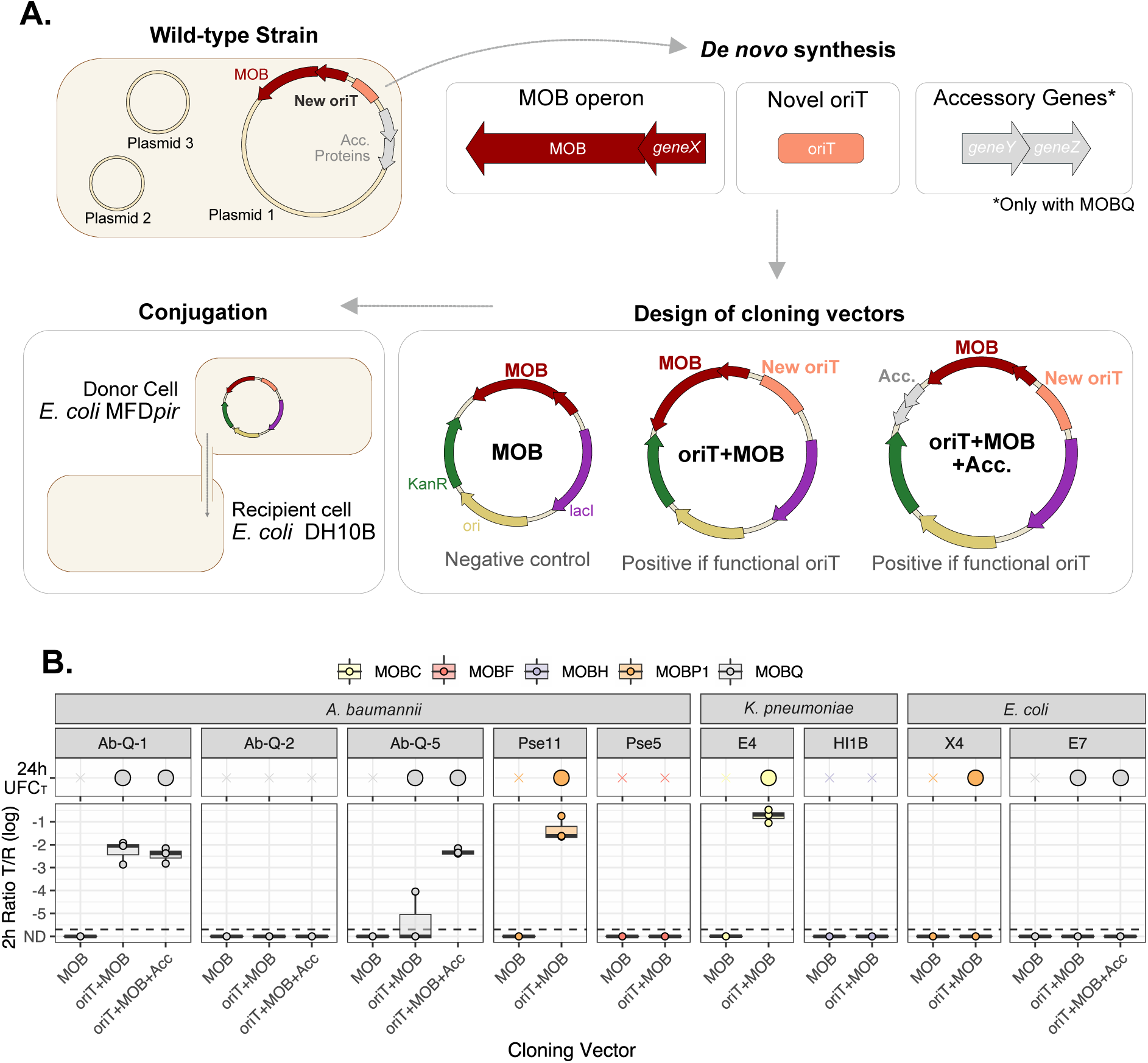
**A.** Schematic summary of the method to validate the novel *oriTs*. The novel *oriT*, relaxase operon, and accessory genes, of natural plasmids were synthesized *de novo* and cloned into a non-conjugative vector. The functionality of the new vector was tested in conjugation assays using as a donor *E. coli* MFD*pir*. **B.** Results of the conjugation experiments. Each facet represents a novel *oriT* tested. The upper plot represents the plasmids in which at least one transconjugant was recovered after 24h (see Methods). In the lower plot, the ratio transconjugants per recipient (T/R) in 2h is represented in a logarithmic scale. The dashed grey line represents the experimental limit of detection (point from which we did not obtained any transconjugant experimentally). ND stands for non-detected, *i.e.*, no transconjugants obtained. The color represents the MOB associated to the *oriT*, being the legend at the top of the figure.

Some of the plasmids showed more conjugation efficiency than others (Table S1). We therefore made 2h mating experiments and observed significant conjugation rates in 4 cases (Figure 5B). Hence, the RP4-conjugation machinery seems to mobilize some of these plasmids at relatively high frequencies. Interestingly, this is the case for three *oriT*s from the distantly related *A. baumannii* species. The three *oriT* that failed to transfer in both types of matings were *oriT*_Ab-Q-2_, *oriT*_HI1B_ and *oriT*_Pse5_. Of note, *oriT*_Pse5_ is present in an *Acinetobacter* plasmid that was shown to be conjugative (pS32-2, MG954379.1)^19^, and our identified *oriT* was found in the previously proposed region (33,345-33,924 nt). Finally, it’s interesting to note than in this heterologous system the incorporation of the accessory genes in MOB_Q_-associated *oriTs* improves conjugation rates in some cases (*oriT*_Ab-MOBQ-5_), but not in others (*oriT*_Ab-MOBQ-1_) (Figure 5B).

### Conjugation mobilizes most plasmids, many carrying antibiotic resistances

To understand the relevance of the 21 novel families of putative *oriTs*, we searched for their occurrences across all available 38,057 plasmids (Table S2). We identified 3,072 new putative *oriTs* in 2,975 plasmids, the vast majority (99%) in replicons of their corresponding families (*Enterobacteriaceae* and *Moraxellaceae*, respectively) (Figure S9). We identified only one putative *oriT* in most plasmid (97%) and only when they lacked a previously known one (98%) (Figure S19). The same way that relaxases are highly associated to a certain Plasmid Taxonomic Unit (PTU)^2^, 12 of the 21 new *oriTs* are in plasmids of a specific PTU (Figure S10). We relied on these results to name the new families of putative *oriTs* after the PTU of the plasmids where they were predominately found (e.g. the new family found in PTU-Pse5 was named *oriT*_Pse5_). The remaining 9 *oriTs* were identified in few plasmids and they lacked an assigned PTU. They were named according to their species and MOB type (e.g. *oriT*_Ec-MOBQ_). With the previous and the novel *oriT* families, we can now identify their occurrences in more than 80% of the replicons of the three species where we presume there should be one (pCONJ, pdCONJ, pMOB) (Figure 6A). The improvement was particularly remarkable in *K. pneumoniae* and *A. baumannii*, where the frequency of pCONJ and pMOB with a putative *oriT* increased from ∼30% and ∼1% to close to 80%. Moreover, our screening identified putative novel *oriTs* in 333 pMOBless, further increasing the fraction of plasmids with a putative mechanism of mobility in these species (72% via conjugation, 7% are phage plasmids) (Figure 6B).

**Figure 6.**
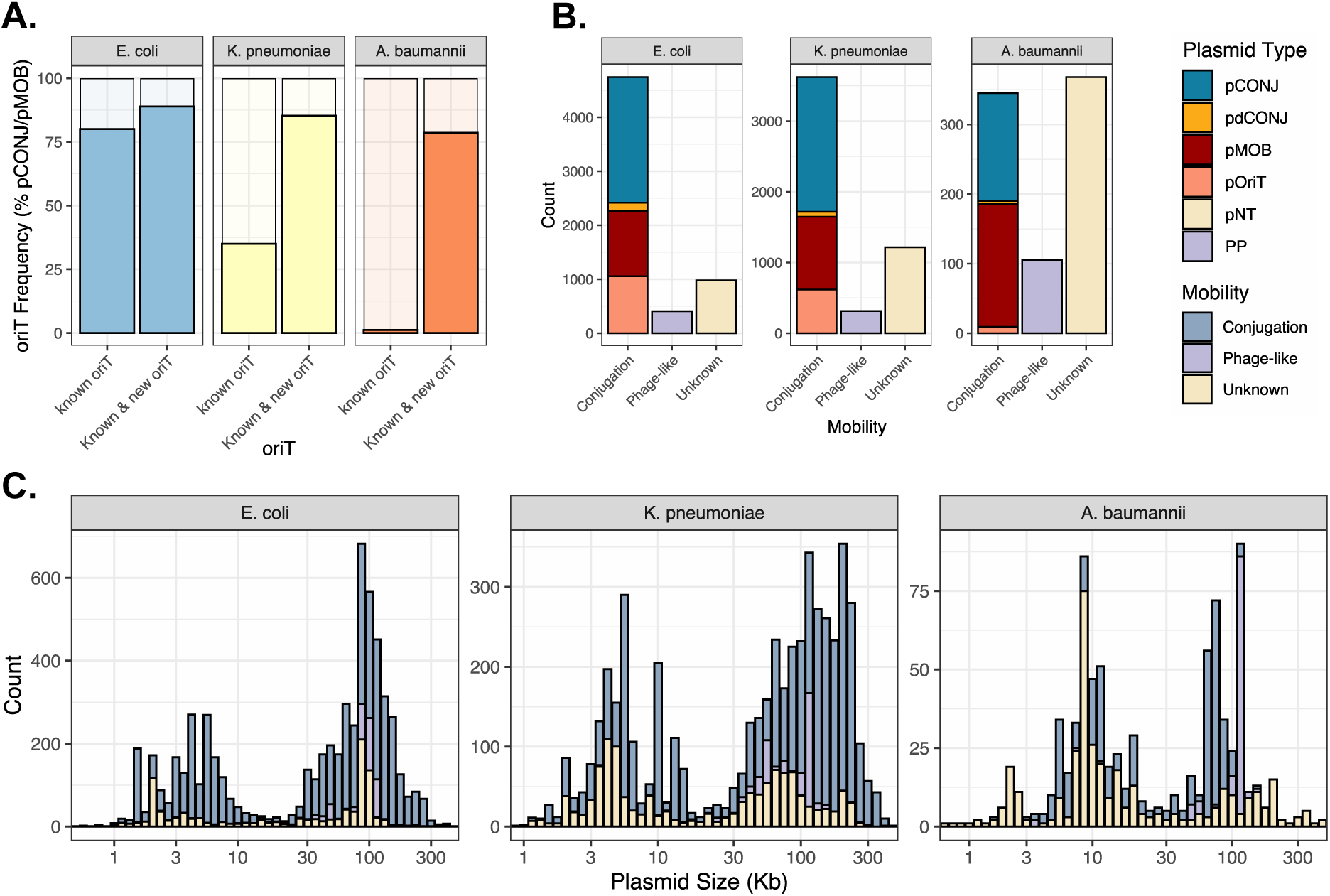
**A.** Frequency of pCONJ and pMOB plasmids carrying *oriTs* in *E. coli*, *K. pneumoniae* and A*. baumannii* screening for the known *oriTs* and with both the known and the new families inferred in our analysis. **B.** Number of plasmids that could be assigned a conjugative mechanism of mobility (pCONJ, pdCONJ, pMOB and pOriT), mobility as phages (PPs), and those plasmids with no known mechanism of transfer (pNTs). **C.** Size distribution of plasmids mobilized by conjugation, like phages, and the presumably non-transmissible.

Plasmids are often carriers of antibiotic resistance genes in nosocomial pathogens, of which *E. coli*, *K. pneumoniae* and *A. baumannii* are among those of highest priority in the clinic^31^. We searched these plasmids to antibiotic resistance genes to assess the potential impact of the findings on novel *oriT*s for the spread of antibiotic resistance. Indeed, some families of plasmids with the novel *oriT*s encode many such genes (Figure S11), some carrying resistance genes to critical antimicrobials (e.g. *oriT*_X4_ associated to *mcr-1*) or a very large number of different resistances (like *oriT*_HI1B_ and *oriT*_FK_). Among the eight novel *oriTs* of *Acinetobacter*, six are associated to the carbapenemase *bla*_OXA_ (Figure S11).

In the end, 15% of *E. coli*, 23% of *K. pneumoniae* and 45% of *A. baumannii* plasmids still lack a recognizable mechanism of mobility (pNTs). These plasmids are found in both peaks of the bimodal distribution of log10-transformed sizes (Figure 6C). Interestingly, 20% of pNTs belong to PTUs where most of the plasmids are either pCONJ or pMOB (Figure S12). They could result from the latter by gene loss^25,32^. In *E. coli* the few remaining pNTs cover a narrow size distribution (∼2 Kb and ∼90Kb), which suggests the presence of a few families of abundant non-transmissible plasmids. About a fourth are related to the known non-conjugative pO157 (∼90Kb, PTU-E5, Figure S12)^33^. In *Klebsiella,* 15% of pNTs are large IncR plasmids (75Kb PTU-R, 56Kb E56, and 61Kb E54), responsible for genetic exchange via conduction, *i.e.*, by co-integrating with conjugative plasmids^34^. We also identified families of small pNTs lacking evidence of mobility in the literature, e.g. the 2Kb Col(BS512) in *E. coli* (PTU-E47, 10% of pNTs) and 4Kb Col440I in *K. pneumoniae* (E71, 7%). The high frequency of the remaining pNTs in *A. baumannii* is intriguing because most pCONJ and pMOB of the species can now be assigned a putative *oriT* (Figure 6A). In *A. baumannii*, 48% of pNT are from three plasmid families: 156Kb PTU-Pse9, 13Kb PTU-Pse7 and the 8Kb PTU-Pse6 (Figure S12). Crucially, conjugation of big plasmids from PTU-Pse9 (e.g. pA297-3 or pD46-4) and >250Kb mega-plasmids (group III-4a, no PTU assigned) has been experimentally confirmed, even if their relaxase (and therefore *oriT*) remain unidentified^35–37^. We tested the hypothesis that conjugative systems of ICEs could be responsible for the mobility of the *A. baumannii* plasmids, as identified in species of *Salmonella* and *Bacillus*^18,38^, but we rarely identified these MGEs in this species (only in 0.5% of the genomes). Hence, the mobility of a few large plasmid families lacking relaxases and whether they are mobilized by conjugation or other means remains to be understood.

## Discussion

Many novel aspects of conjugation have been uncovered in the last few years, including the magnitude of *in trans* conjugation^39^, the mobility through RC-Rep proteins^18,40^, the stabilization of conjugative pili by cell receptors^41^, the mechanisms of ssDNA to dsDNA conversion after plasmid acquisition^42^, the cost of mobility^43,44^, and the anti-defense arsenal encoded in conjugative leading regions^45^. Systematic large-scale analyses of plasmids have broadened our understanding of these processes^17,22,23,45,46^. Yet, we still ignore a lot about conjugation across most of the bacterial realm because we ignore how many plasmids are mobilizable. Here, we used previous data to characterize known *oriTs* and use this information to establish a method to identify novel ones. This uncovered 21 new putative families of *oriTs* in *E. coli*, *K. pneumoniae*, and *A. baumannii*. This approach can be extended to other bacterial taxa and spur a renewal of interest on types of conjugative systems that have not been sufficiently studied. Being able to identify *oriTs* will also facilitate genetic engineering in non-model bacteria.

There is usually strong genetic linkage between DNA motifs and their interacting proteins in MGEs. When recombination is frequent, as in plasmids^34,47^, this protects the functional module from being split into different mobile genetic elements resulting in loss of function. We show that *oriTs* are systematically close to the relaxases. Most exceptions are caused by linkage with other conjugation genes (*oriT*_pVCR94Λ1X_^9,48^) sometimes in the operon of the relaxase (*oriT*_F_). The only clear exception is *oriT*_pAD1-H_, firstly described within the *repA* gene of the plasmid pAD1^49^. This *oriT* is atypical: it is often within coding sequences (Figure S3) and found far from identifiable conjugation genes (Figure 2E). The novel *oriTs* are always close to the relaxases and intergenic (Figure S10). This is not a strictly necessary consequence of our method to identify them, since we only require that a sizeable fraction of the *oriTs* of a family are intergenic and close to the relaxase. The strong genetic linkage between conjugation genes and *oriT* and its position in intergenic sequences can thus be used to narrow the space of search for novel *oriT*s.

Occurrences of the *oriTs* of the same family are expected to be recognized by relaxases of the same type and the inverse is often also true. Most relaxases belong to the superfamily of HUH endonucleases, which tend to recognize and cleave DNA motifs in non-coding regions^50^. This location may facilitate the evolution of *oriT*s, which would be more constrained if they overlapped coding sequences. It may also facilitate the formation of secondary structures to improve DNA-relaxase interactions (that could be disrupted by or disrupt the transcription machinery)^51^. In spite of their intergenic location and the often-rapid evolution of HUH endonucleases DNA targets, the conservation of *oriTs* is quite high. When relaxases are more than 60% identical in amino acid sequence they are almost always associated with a recognizable *oriT* of the same family. This suggests tight co-evolution between the relaxase and the cognate *oriT*. It also suggests that we can extend our method to uncover *oriTs* across most bacterial conjugation systems for which we know the relaxases.

The quality of our predictions of novel *oriT* families is further supported by additional characteristics not used in the method to identify them. We showed that *oriTs* are associated with specific PTUs, specific relaxases, and specific bacterial hosts. They tend to be associated with either conjugative plasmids or plasmids with relaxases (pMOB), but not both. They tend to be surrounded by secondary structures and often have *nic* sites known for the cognate type of relaxases. Nevertheless, our approach cannot yet precisely delimit the important regions of the *oriT*. When analyzing the literature, we could not find a consensus, with *oriT* definitions varying from the minimal region needed for mobilization (e.g. 31 nt for R64)^52^, to the large (∼350 nt) region including the binding of various relaxosome-accessory proteins^54^. Beyond a problem of definition, computational delimitation of *oriTs* is difficult because it requires to separate the similarity caused by natural selection of the *oriT* sequence from the similarity caused by phylogenetic inertia and by other DNA motifs, such as promotors or origins of replication. Further work will be necessary to uniformly characterize the conserved functional positions of the novel (and many of the ancient) *oriT*s.

We have confirmed that 6 out of 9 tested *oriTs* allow the mobility of plasmids *in trans* by the RP4 conjugation system in *E. coli*. Two of the three *oriT* that failed to transfer, *oriT*_HI1B_ or *oriT*_Pse5_, are from *oriT*s of conjugative plasmids. We previously discussed that relaxases of conjugative systems may be less promiscuous in their interactions with the conjugation systems than those of mobilizable plasmids^25,55^. This may explain why they failed to work in a system that is very distinct from the natural one. Still, the experimental data provides a clear proof of principle for our method and shows that we successfully identified functional origins of transfer in plasmids. Beyond plasmid biology, this is important to understand the spread of antimicrobial resistance. The tested *oriT*s are found in many plasmids carrying some of the most worrisome antimicrobial resistance genes in three species that are themselves among those posing the higher risk in clinical settings^31^. This may facilitate a better understanding of their epidemiological patterns of occurrence.

We found putative *oriTs* in most of the relaxase-encoding plasmids. We also identified hundreds of novel pOriTs. Yet, some plasmids still resist classification in terms of transmissibility. In *E. coli* and *K. pneumoniae*, many of them could be assigned to specific well-known non-conjugative plasmids. The case of *Acinetobacter* is particularly intriguing since half of the plasmids lacking a relaxase also lack *oriTs* like those of the other plasmids and ICEs. Some big plasmids might encode still uncharacterized relaxases and MPF systems^35,37^. Nevertheless, most of these plasmids are very small (Figure 6B), and they may be mobilized by natural transformation in *A. baumannii* as recently observed for small cyanobacteria plasmids^56^. In contrast, in *E. coli* and *K. pneumoniae* our results suggest we are not far from having identified the full set of potentially mobile plasmids in the extant genomes. The presence of origins of transfer across so many plasmids establishes an intricate network of dependencies between them that is now open for exploration.

## MATERIALS AND METHODS

### Genome data

We retrieved all the complete genomes from bacteria available in the NCBI non-redundant RefSeq database (https://ftp.ncbi.nlm.nih.gov/genomes/refseq/, last accessed in May 2023). This resulted in a set of 32,799 genomes. Only the replicons tagged as plasmids were retrieved for further analysis resulting in 38,057 plasmids (Table S2).

### Plasmid classification

Plasmids were classified according to the different schemes available: (1) Plasmids were assigned to a Plasmid Taxonomic Unit (PTU) when possible using COPLA, version 1.0^57^; (2) Each plasmid was assigned to a Incompatibility (Inc) group according to PlasmidFinder, version 2.0.1 (database available on January 2023)^58^; (3) Plasmids encoding a relaxase and/or a mating pair formation (MPF) system were assigned to a specific MOB group and MPF type by the module CONJscan of MacSyFinder, version 2.0^59^.

### Functional annotation of the plasmid database

The plasmids were annotated using Prodigal v2.6.3^60^, with the option *-p meta* recommended for mobile genetic elements. The genes related to antimicrobial resistance, virulence and stress response were identified with AMRFinderPlus, version 3.11.4^61^. We used the module CONJscan of MacSyFinder, version 2.0^59^, with the models specific for Plasmids, to identify protein coding genes related to conjugation, such as relaxases and the MPF system. CONJScan was also used to screen for putative ICEs (complete MPF systems) and IMEs (only MOBs) in the chromosomes of *E. coli*, *K. pneumoniae* and *A. baumannii*. Phage-like functions characteristic of Phage-Plasmids were identified using the pipeline recently described^17^.

### Identification of known origins of transfer

We updated a previously published collection of 91 experimentally validated *oriTs*^22,62^, including two more *oriTs* from the literature^63,64^. Given the high sequence similarity between some *oriTs*, e.g. 100% identity between *oriT*_R6K_ and *oriT*_pMAS2027_ or 86.2% identity between *oriT*_F_ and *oriT*_pSU233_, we clustered them by sequence similarity to obtain *oriT* families.

Briefly, an ‘all-vs-all’ local alignment of the *oriTs* was performed using the EMBL-EBI EMBOSS Needle Smith-Waterman algorithm, version u’2022-09-13 12:15’^65^. We retrieved the hits with >70% identity and >70% coverage (of the smaller sequence). This threshold was selected since, below this parameters, unrelated *oriTs* are grouped in the same family, like the *oriT*_ColE1_ (characteristic of MOB_P5_-encoding pMOBs) and the *oriT*_R64_ (MOB_P12_-encoding pCONJ)^66^. The hierarchical clustering was performed using the Ward.D2 algorithm of the R package pheatmap, version 1.0.12 (https://CRAN.R-project.org/package=pheatmap). Among the 93 *oriTs*, 43 *oriTs* were grouped into 13 clusters. Each of these clusters was subsequently inspected to detect anomalous clustering: the *oriT* sequences of each cluster were extracted, a multisequence alignment (MSA) was performed using mafft, version v7.505^67^, with the options *--maxiterate 1000 --genafpair*. The output was expertly curated using the visualization software SeaView 5.0.4^68^. This led to the exclusion of *oriT*_ICEKpl_, *oriT*_Tn1549_, and *oriT*_CloDF13_ from their respective clusters, because they revealed very poor sequence alignments with the remaining sequences. At the end of this process, we obtained 40 *oriTs* clustered in 13 *oriT* families, plus 53 *oriT* singletons (*i.e.*, not clustering with other sequences, Figure S13).

To identify the *oriTs* in our plasmid collection, we used the BLAST suite of programs, version 2.9.0+^69^. We indexed the plasmid collection using makeblastdb (default options). Then, we used blastn, options *-task blastn-short -evalue 0.01*, to screen for the *oriTs* (query) against the 38,057 plasmids (database). Only the hits with an identity >80% and coverage >80% were retrieved. Exceptionally, we adapted the criteria for two *oriT* queries that are particularly short (13 bp, *oriT*_CloDF13_ and *oriT*_pAD1_), in which case only 1 mismatch was allowed. When two different types of *oriT* were identified in the same region of a plasmid (overlapping), we assigned the *oriT* hit to the type with the best E-value. For simplification, when a hit corresponded to a member of an *oriT* family, we maintained the family name (e.g. if we identified the *oriT*_pKL1_ in a plasmid, we used the family name “*oriT*_R64_” for the following analysis because the two *oriTs* belong to the same family) (Figure S13). Exceptionally, despite *oriT*_pAD1_ being originally associated to MOB_C_, we identified a similar sequence often associated to MOB_H_. We therefore referred to this *oriT* as *oriT*_pAD1-H_. We identified an exceptional case of a 54kb *E. coli* plasmid with 23 occurrences of an *oriT*_ColE1_. This plasmid (NZ_CP019265.1) was discarded from further analysis as we suspect it is a sequencing artifact.

### Association between the MOB and *oriT*

To assess how similar are the relaxases associated to the same *oriT* families, we analyzed the relaxases identified by CONJScan. Among the 20,953 relaxases identified in 19,745 plasmids, we excluded those coming from plasmids encoding multiple-relaxases and/or carrying multiple *oriTs*. Exceptionally, plasmids encoding a MOB_P1_ relaxase with x2 *oriT*_R6K_ were also retrieved for the analysis since these *oriTs* are typically duplicated in the plasmids^70^. This resulted in a total number of 18,439 relaxases. Some relaxases are very similar. To account for this, we clustered very similar relaxases (99% identity in protein sequence, 99% coverage) with MMseqs2, version 13.4511, and picked one protein per cluster. This resulted in a set of 6,547 relaxases, which were split into the different MOB groups. We then compared all-vs-all relaxases within each MOB group using blastp, version 2.9.0+ ^69^, and retrieved the best bidirectional hits (BBH) when the E-value was <0.01 and the alignment covered at last 50% of the smaller protein.

### Phylogenetic analysis of the relaxases

We built the phylogenies of the 7,711 relaxases of *Escherichia coli*, *Klebsiella pneumoniae*, and *Acinetobacter baumannii* by previously splitting them into different MOB groups and then building a tree for each group. To do so, we built an MSA using mafft, version v7.505^67^, options *--maxiterate 1000 --genafpair*. The MSAs were then trimmed using the software ClipKIT, version 1.3.0^72^, option *--gappy*. The phylogenetic tree was built using IQ-Tree, version 2.0.3^73^, with the ultrafast bootstrap option *-B 1000*^74^ and the best model finder *-MFP*^75^. For the visualization of the trees and their metadata, we used the online tool iTOL^76^.

### Genetic localization of the origins of transfer

To analyze the location of the *oriTs* relative to coding sequences (CDS) (*oriT* within a CDS vs. *oriT* within a non-coding sequence), we analysed their position relative to the functional annotation file. Origins of transfer whose sequence was partially identified within a gene, were considered genic when >80% of their sequence was included in the CDS. To analyse the location of the *oriTs* relative to the relaxase, only plasmids with 1 MOB and 1 *oriT* were considered (and exceptionally 1 MOB_P1_ with 2 *oriT*_R6K_). Additionally, only plasmids with the canonical MOB-*oriT* pair were analysed. We defined the canonical MOB-*oriT* pair as the most frequent MOB type identified for a given *oriT* family. To assess the position of the *oriTs* relative to the relaxase (upstream/downstream), we used the shortest circular distance between the *oriT* and the relaxase in every pCONJ, pdCONJ and pMOB. If the *oriT* was closer to the 5’ end of the gene than to the 3’, the *oriT* was considered to be upstream to the MOB, and vice-versa. Likewise, the distance between the *oriT* and every conjugative gene of the MPF system of pCONJ was calculated. The later was visualized using the R package gggenes, version 0.4.1 (https://CRAN.R-project.org/package=gggenes), setting the 5’ of the *oriT* as position 0 bp.

### Computational approach to identify novel origins of transfer

To infer new *oriTs*, we retrieved the first intergenic sequence upstream the 20,953 MOBs identified among the 38,057 plasmids. If the first intergenic sequence is smaller than 50bp then it was not considered, and we took the second one (and so on). Intriguingly, 10 plasmids did not have any such intergenic sequence (presumably due to sequencing artifacts) and were discarded from the analysis. Therefore, 20,943 non-coding sequences ≥50 bp were used. Redundant sequences were identified by clustering the DNA sequences by sequence similarity (99% identity, 99% coverage) with MMSeqs2, version 13.451^71^, and one of each cluster was picked randomly for subsequent analysis. This resulted in a dataset of 7,955 sequences. We calculated an ‘all-vs-all’ local alignment of the non-coding sequences associated with protein coding genes within each MOB family. This was done using the optimal Smith-Waterman algorithm (*ssearch36*) of the FASTA package^77^. We only retrieved hits with an E-value <0.001, identity >60%, and an alignment length >50 nucleotides. Every sequence with no hits but to itself was discarded. The bitscore of the retrieved hits were then normalized (bs_A-B_+bs_B-A_)/(bs_A-A_+bs_B-B_) and used for a hierarchical clustering analysis of the non-coding sequences using the Ward.D2 method of the R package pheatmap, version, version 1.0.12.

The clustering to infer new *oriTs* in *E. coli*, *K. pneumoniae* and *A. baumannii* was performed in 3 rounds to successively discard large clusters with already known *oriTs* (Figure S14). In each of the rounds, 30 clusters were built, and those including known *oriTs* (or present in plasmids with known *oriTs* far from the relaxase, e.g. *oriT*_F_, *oriT*_pVCR94X_, *oriT*_pAD1-H_) were discarded for the following round. This way, among the 1,061 initial sequences, we retrieved 60 clusters: 36 clusters (577 sequences) carrying known *oriTs*, and 24 clusters (232 sequences) with novel putative *oriTs*. Additionally, 252 sequences were singletons or in small groups and were not further investigated. For each of the clusters we made a MSA of their sequences using mafft *--maxiterate 1000 --genafpair*, version v7.505^67^. The quality of the MSA was carefully examined with the software SeaView 5.0.4^68^. To note, the *oriT*_R64_, *oriT*_ColE1_, and novel putative *oriT*_E7_ were split into 2, 6 and 3 redundant clusters respectively, so their sequences were grouped together. Finally, a cluster belonging from upstream regions of *Klebsiella* MOB_F_ was discarded, since it corresponds to a non-coding sequence sometimes present upstream the putative *oriT*_FK_. The non-coding sequences carrying a new putative *oriT* are available in File S1.

### Computational examination of the novel putative origins of transfer

Each cluster of non-coding sequences was examined for the presence of already known *oriTs* (using blastn as previously described). Additionally, we checked the size of their sequences, their distance to the relaxase, their secondary structure, the presence of known *nic* sites of *oriTs*, and the location of their corresponding relaxases in a phylogenetic tree of the relaxases of each species (built as previously described). Lastly, one representative per cluster was selected, and screened in the plasmid database using blastn, E-value <0.01, identity >70%, and coverage >70%. We then analyzed the similarity between relaxases associated with *oriTs* of the same family (as mentioned above for known previously *oriTs*). An ‘all-vs-all’ alignment of the relaxases protein sequences was done using blastp, version 2.9.0+^69^, and retrieved the best bidirectional hits (BBH) when the E-value is <0.01 and the alignment covered at least 50% of the smaller protein.

### Secondary structure of the origins of transfer

To infer the secondary structure of the known *oriTs* and non-coding regions bearing the putative new *oriTs*, we employed the RNAfold web server of the Vienna RNA Websuite tools, with the default folding algorithms of minimum free energy with the DNA parameters^78^.

### Cloning vectors used for the experimental validation

To validate the novel *oriTs*, we selected nine candidates from different species (*E. coli*, *K. pneumoniae*, *A. baumannii*), MOB types (C, F, H, P1, Q), and mobility categories (pCONJ, pMOB) (Table S1). The novel *oriTs* and relaxase operons were synthesized *de novo* and cloned into pCOLADuet^TM^-1 by GenScript USA Inc. (Figure S9). As a negative control, each of the relaxases was independently cloned into pCOLADuet^TM^-1 without the *oriT* (Figure S9). The empty pCOLADuet^TM^-1 was also used as a negative control. In MOB_Q_ plasmids, some genes upstream the *oriT* have a role in the relaxosome formation^29^. In these cases, a third vector was obtained, with the one/two gene/s upstream the *oriT* cloned into pCOLADuet^TM^-1 in addition to the putative *oriT* and MOB (Figure S9). Hence, for each *oriT*, we used the pCOLADUET^TM^-1 with the MOB only, MOB+*oriT*, and MOB+*oriT* plus accessory genes in MOB_Q_ cases. Details on the plasmids is available in Table S1 and their DNA sequence in File S4.

### Conjugation assays

We used *E. coli* MFD*pir* as donor strain and *E. coli* DH10B as recipient (Table S2). *E. coli* MFD*pir* carries the conjugative machinery of RP4 in the chromosome^24^, potentially being able to mobilize the pCOLADuet^TM^-1 derivatives. Both donors and recipients were independently grown overnight in 5ml Lysogeny Broth (LB) at 37°C. Donors were supplemented with 0,3mM diaminopimelic acid (DAP) (auxotrophic for this amino acid) and 50μg/ml of Kanamycin (to select for pCOLADuet^TM^-1). Recipients were supplemented with 50μg/ml Streptomycin. The overnight cultures were then diluted 1/50 into fresh media (5ml) and grown under the same conditions until they reached an OD_600_ 0.8-1. Then, donors and recipients were mixed in a ratio 1:1 (400μl each). The mix was centrifuged (2min, 6000g) and the supernatant discarded. The pellet was twice resuspended in 1ml phosphate-buffered saline (PBS) and centrifuged (2min, 6000g). The resulting pellet was resuspended in 10μl PBS and added to agar pads in 24-well plates. Each well contained 1ml of LB with 0,3mM DAP and 1mM isopropyl β-D-1-thiogalactopyranoside (IPTG 10mM) to induce pCOLADuet^TM^-1 promotors. The plate was incubated at 37°C for 2h. After the conjugation time, the agar pad was resuspended in 1ml PBS and incubated for 5min (37°C, 30rpm). The ml was then centrifuged (2min, 6000g), the supernatant discarded, and the pellet resuspended in 50μl PBS. Then, 20μl were transferred to 180μl PBS in a 96-well plate, and serial dilutions were performed (20μl in 180μl) reaching up to 10^−7^. 5μl of each dilution was spotted on plates selecting for transconjugants (Str+Km), transconjugants and donors (DAP+Km), and recipients (Str). CFUs were counted after an overnight (37°C) at the spots with 5-25 CFUs. A single colony would be equivalent to 2×10^3^ CFUs/ml. To recover transconjugants of plasmids beyond the detection limit, we performed the same conjugation assay with an increased mating time (24h) and culturing all bacterial load after conjugation.

## Acknowledgements

We would like to thank Dr. Fernando de la Cruz and Dr. María Pilar Garcillán Barcia for their help and advice on this project. Additionally, we would like to thank Dr. Rémy A. Bonin and Dr. Xavier Charpentier for providing biological material. Lastly, we would like to thank to all the Microbial Evolutionary Genomics Unit for the regular discussions throughout the project. This study was funded by the INCEPTION project (PIA/ANR-16-CONV-0005), the Fondation pour la Recherche Médicale (Equipe FRM, EQU201903007835) and the European Commission (HORIZON-MSCA-2021-PF-01–01 EvoPlas-101062386). This work used the computational and storage services (TARS cluster) provided by the IT department at Institut Pasteur, Paris.

## SUPLEMENTARY INFORMATION

**Table S1.** Information on bacterial and plasmids used in the conjugation experiments.

**Table S2.** Complete information regarding the 38,057 plasmids used in this work. The RefSeq accession number is provided, together with data on the taxonomy, size, PTU, mobility and genes encoded (antimicrobial resistance, virulence and stress response).

**File S1.** Multifasta with the DNA sequence of the intergenic regions belonging to the clusters that represent putative novel *oriTs*. The header of each sequence represents the new *oriT* and the accession umber of the plasmid in which it is found.

**File S2.** Newick format of the phylogenetic tree of the MOB_P1_ relaxases shown in Figure 4B.

**File S3.** Newick format of the phylogenetic tree of the MOB_Q_ relaxases shown in Figure 4C.

**File S4.** Multifasta with the DNA sequences of the cloning vectors used for the conjugation experiments. The header of each sequence indicates the novel *oriT* family tested and the genes inerted into the vector pCOLADuet-1^TM^ to test it.

**Figure S1.**
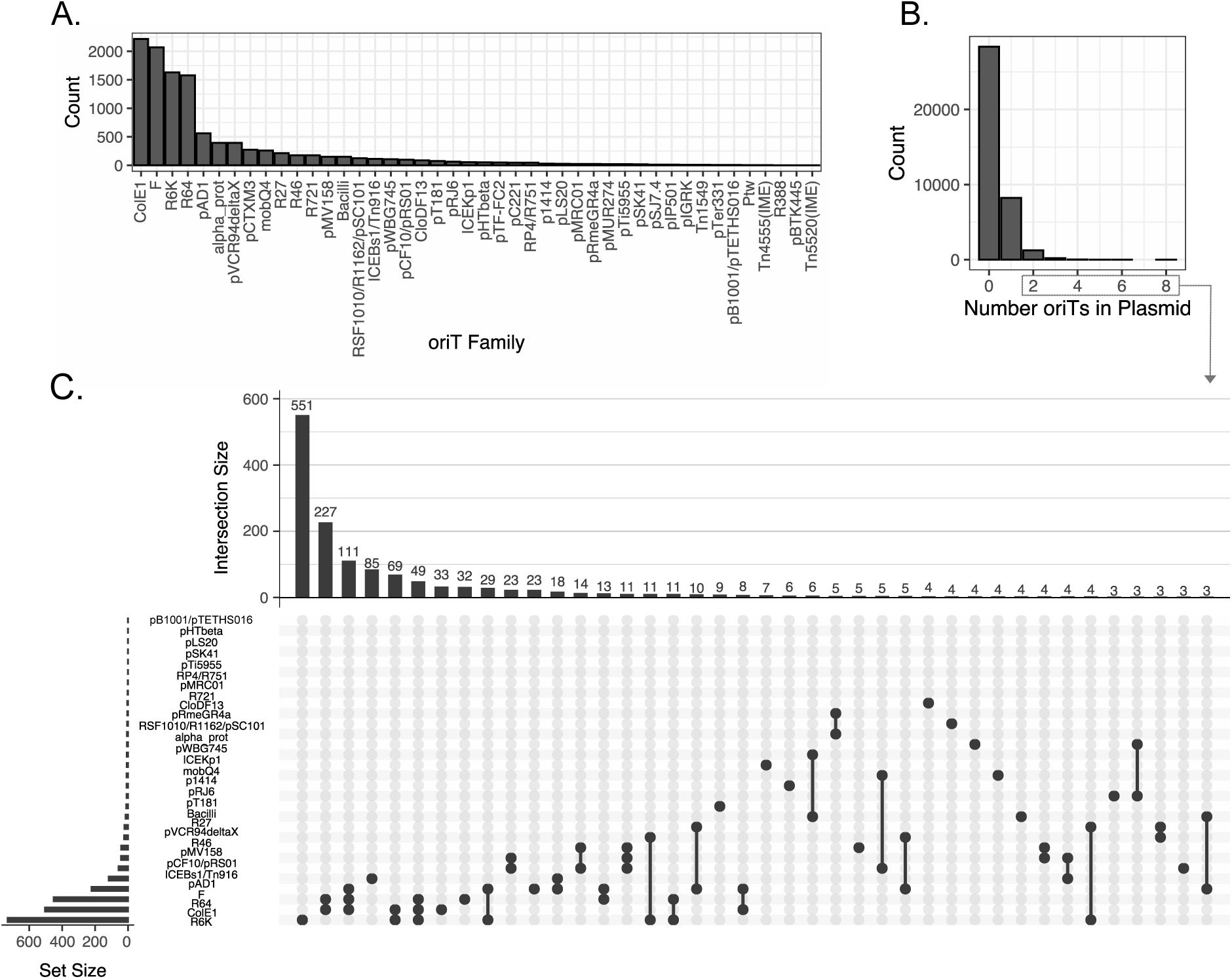
**A.** Families of known origins of transfer identified in all Bacterial plasmids. **B.** Number of *oriTs* identified per plasmid. **C.** Co-occurrence of *oriT* families in the same plasmid. Only plasmids with more than 1 *oriT* are shown. If only one dot is shown, it means that the same *oriT* is found more than once in the plasmid.

**Figure S2.**
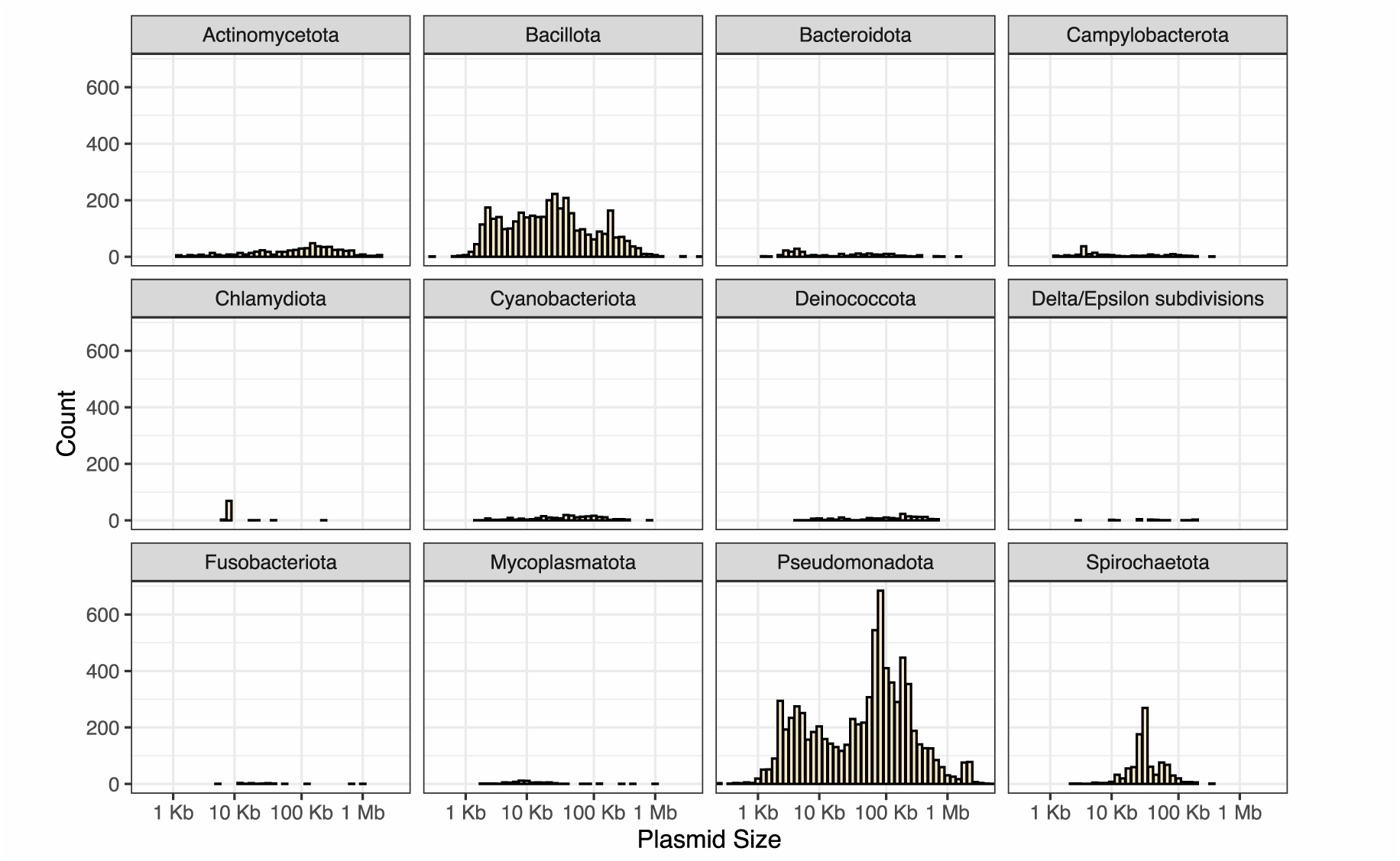
Plasmid size of the plasmids lacking a previously known origin of transfer in phyla with more than 25 plasmids in RefSeq.

**Figure S3.**
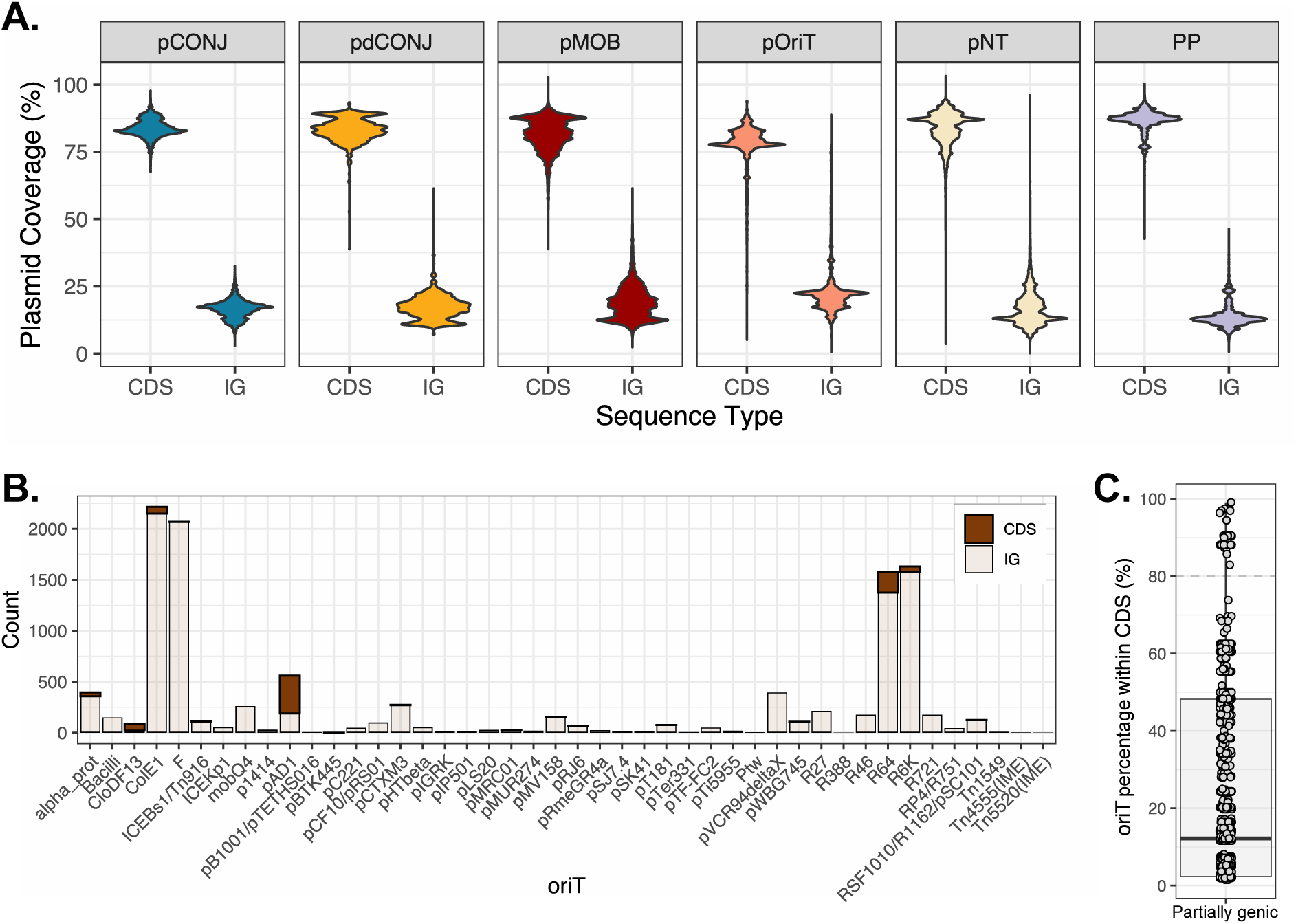
**A.** Fraction of each plasmid sequence that corresponds to coding sequences (CDS) or to intergenic regions (IG) according to its mobility category. **B.** Per *oriT* family, number of *oriTs* identified in coding sequences (CDS) or in intergenic sequences (IG). **C.** Percentage of the *oriT* sequence overlapping with a coding sequence among the 2,633 *oriTs* partially identified within a CDS. The dashed grey line (80%) represents the threshold used to split them into coding (>80%) or intergenic (<80%).

**Figure S4.**
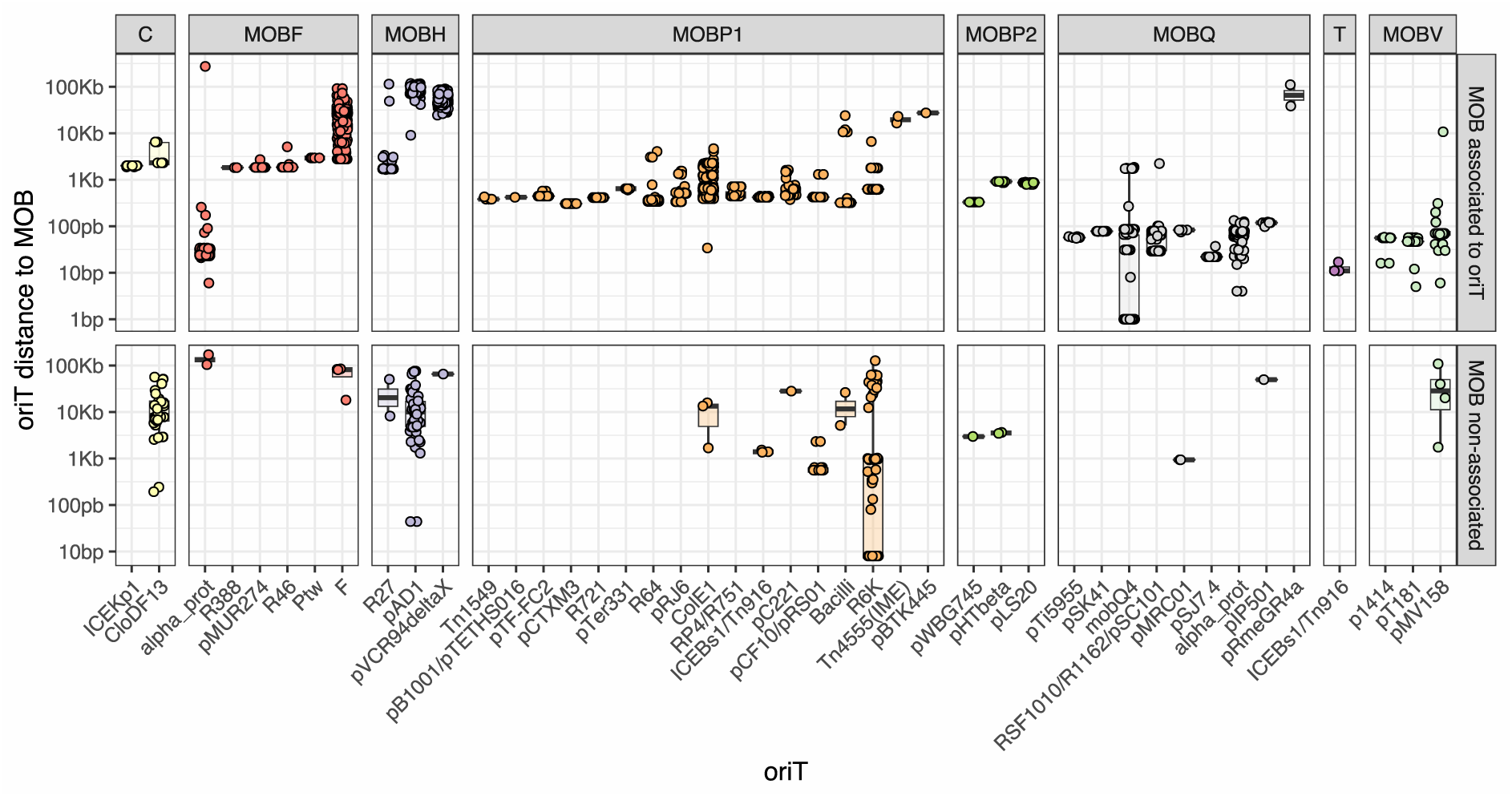
Distance between the *oriT* and the relaxase (MOB). The upper part shows the distance of each *oriT* in plasmids encoding the canonical *oriT*-MOB associations (e.g. MOB_F_ and *oriT*_F_, MOB_P_ and *oriT*_ColE1_). In the lower part of the figure, it is shown the distance of the *oriT* to the only relaxase in the plasmid when the MOB type was not the canonical one for the specific *oriT* (e.g. MOB_P1_ and *oriT*_F_). These non-canonical associations represent 3% of the cases.

**Figure S5.**
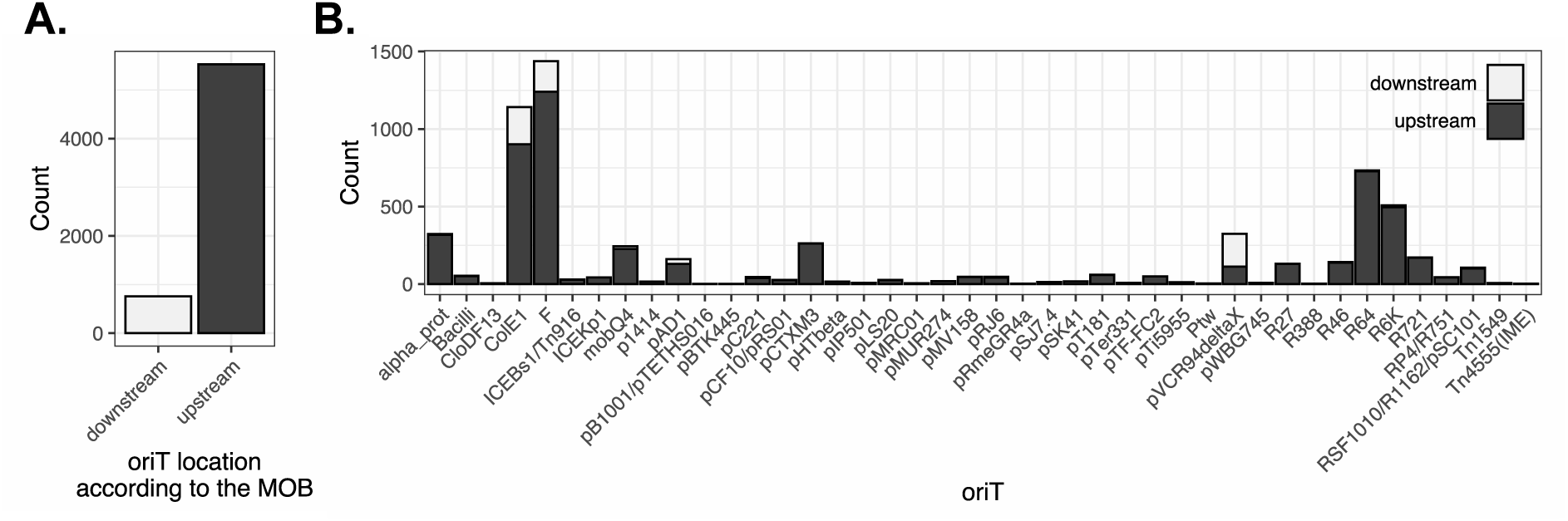
**A.** Position of the *oriT* relative to the relaxase in all the plasmids encoding one relaxase and one *oriT* (or 2 *oriT*_R6K_). **B.** Position of the *oriT* relative to the relaxase per *oriT* family.

**Figure S6.**
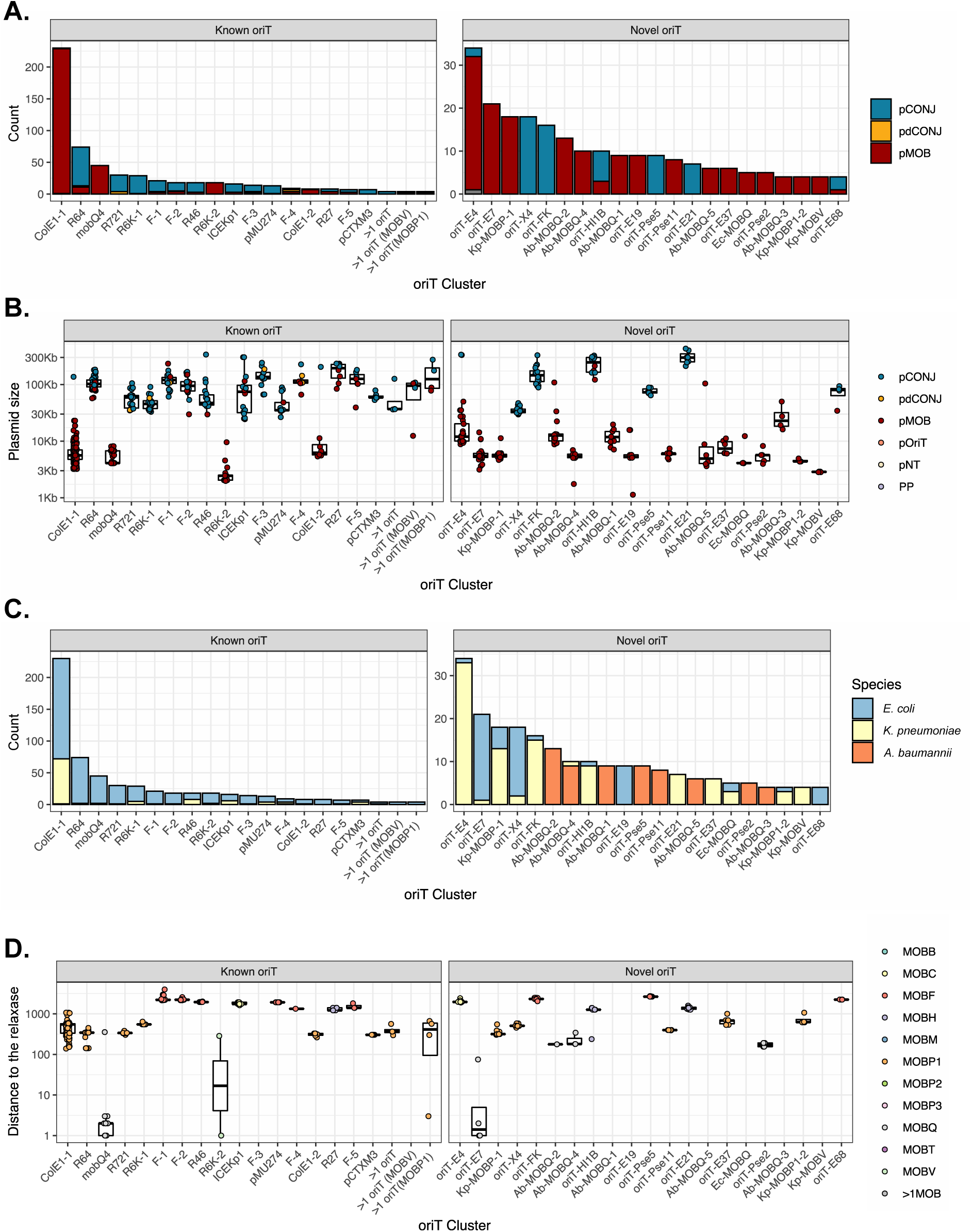
**A.** Mobility type of the plasmids in which the members of the clusters of the new putative *oriT* families are encoded. **B.** Size of the plasmids in which the *oriT* clusters have been identified. **C.** Bacterial species in which the intergenic sequences were identified**. D.** Distance between the new putative *oriT* and the relaxase encoded in the plasmid. The color of the dots represent the MOB type. All the legends are at the right of the figure.

**Figure S7.**
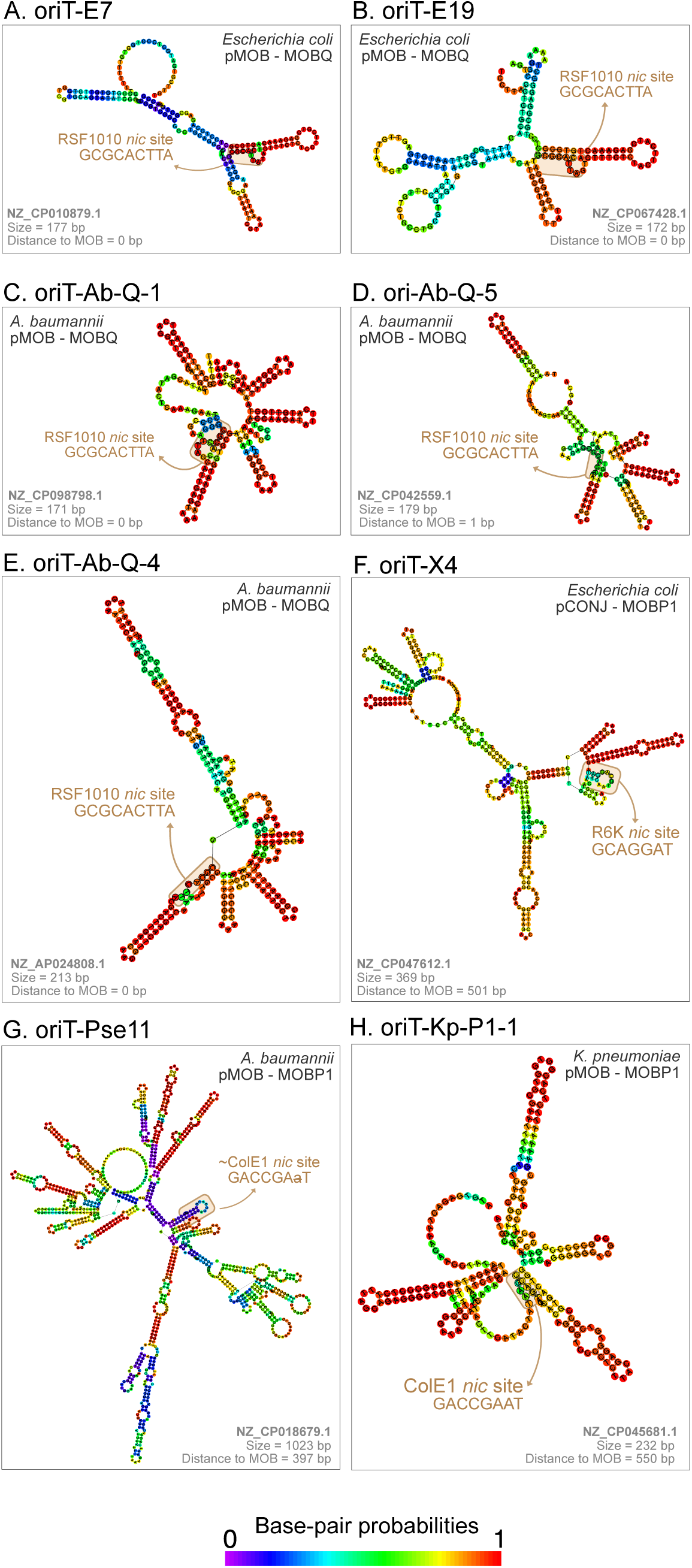
Secondary structure and *nic* sites of putative new *oriT* families. Information on their plasmid accession number, mobility, MOB type, intergenic sequence size, and distance to the relaxase, is shown within the figure.

**Figure S8.**
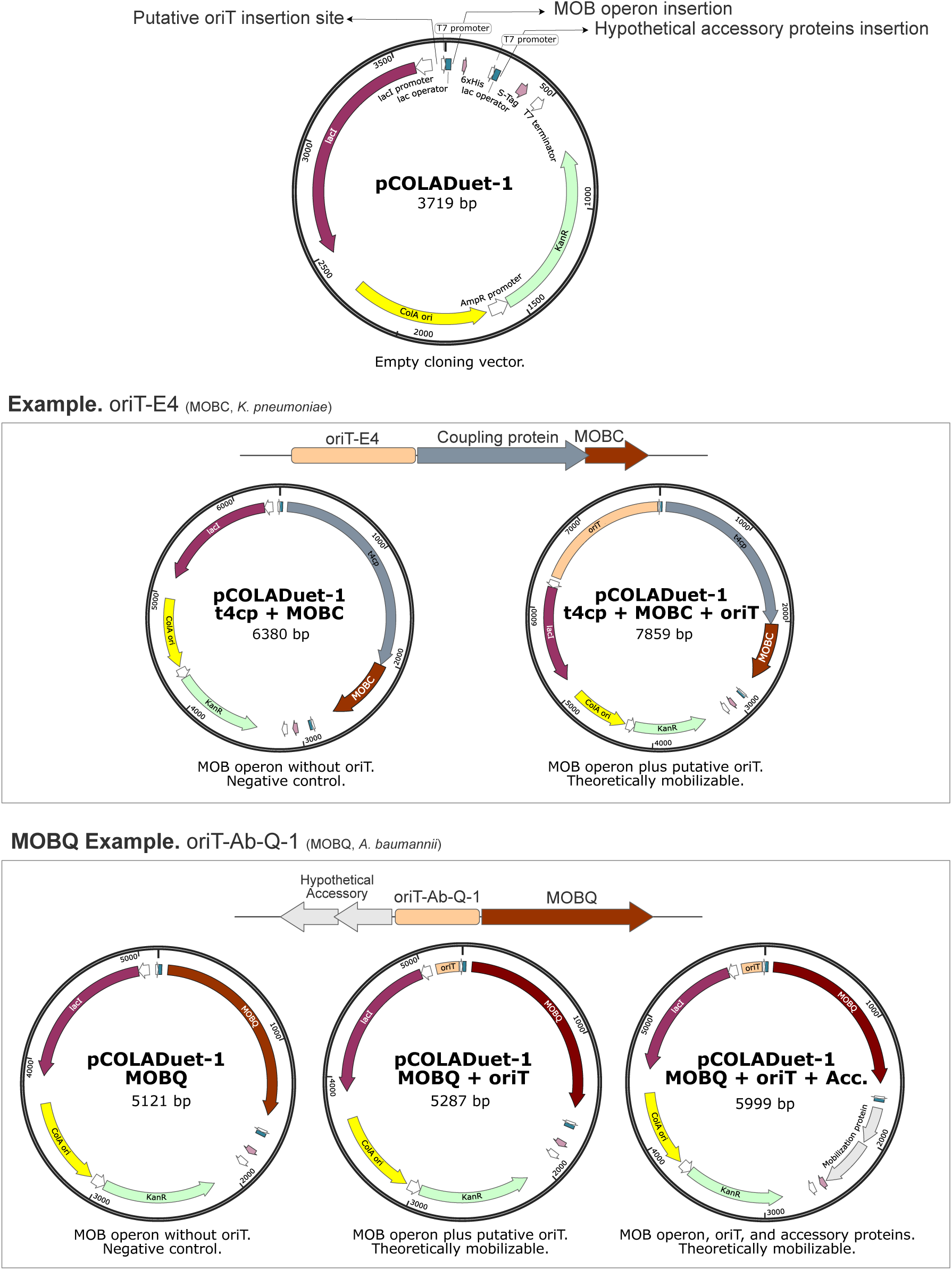
Map of pCOLADuet-1 with the insertion sites to validate the novel *oriTs*. As an example, the wild-type genetic environment of *oriT*-E4 used in the experimental validation is shown. As negative control we designed pCOLADuet-1 with the MOB operon (coupling protein-MOB_C_), while pCOLADuet-1 with the MOB operon and *oriT*_E4_ is used to test the functionality of this *oriT*. As a MOBQ example, the wild-type environment of *oriT*_Ab-Q-1_ is shown. As a negative control, we used pCOLADuet-1 with the MOB operon (only MOB_Q_). To test the *oriT*, two different vectors were used (with and without the accessory proteins opposite to the MOB_Q_). These double constructions were only built for *oriTs* associated to MOBQ relaxases

**Figure S9.**
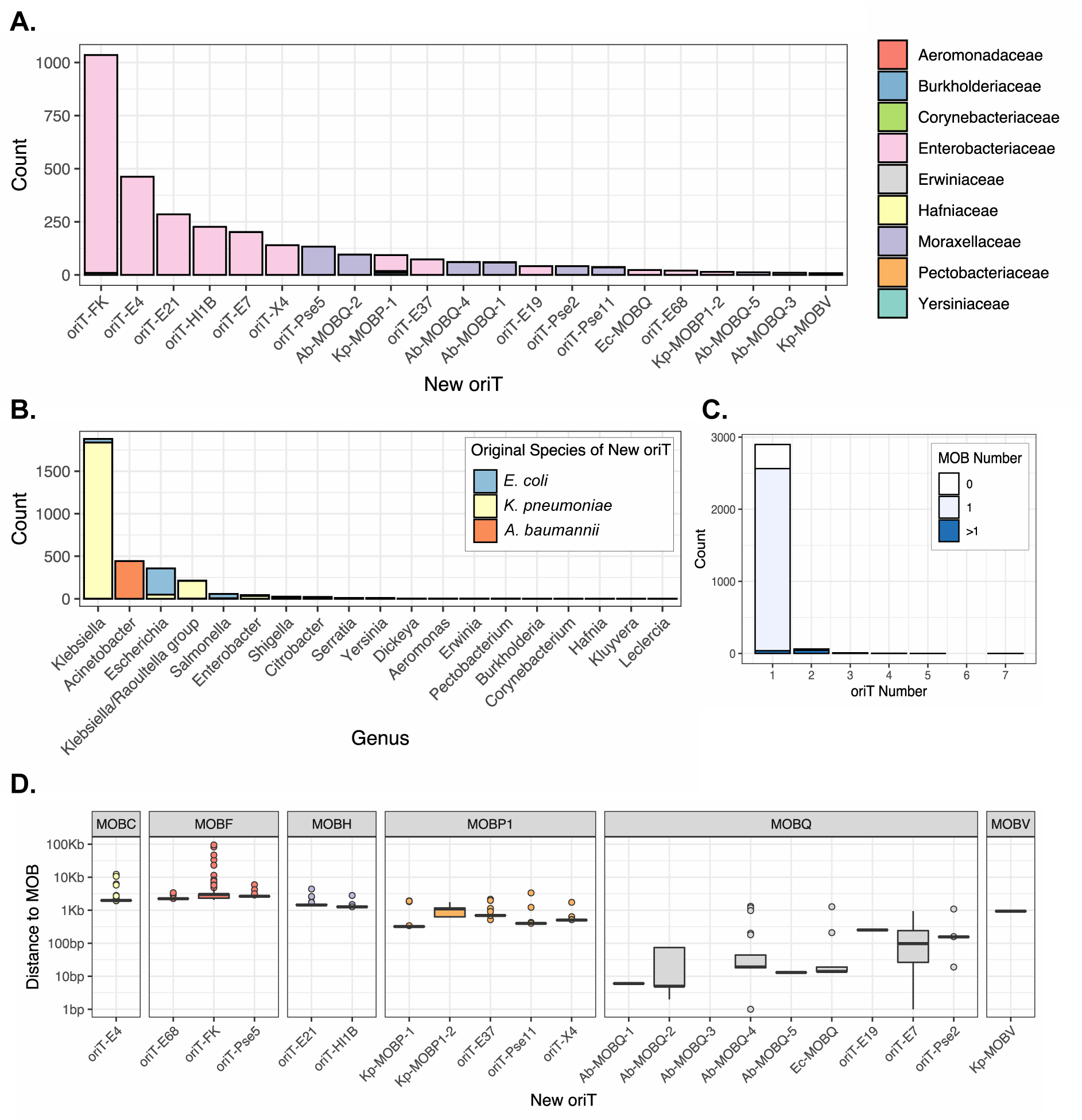
**A.** New putative *oriTs* identified among the 2,975 plasmids. The colors represent the family host of the plasmid carrying the *oriT*. **B.** Genera in which the new putative *oriTs* have been identified. The color indicates the species in which the *oriT* was originally identified in our analysis. **C.** Number of new *oriTs* identified per plasmid. The color shows the number of relaxases in the plasmids. **D.** Distance of the relaxase to the novel *oriTs*. Only plasmids with 1 *oriT* and 1 relaxase were included.

**Figure S10.**
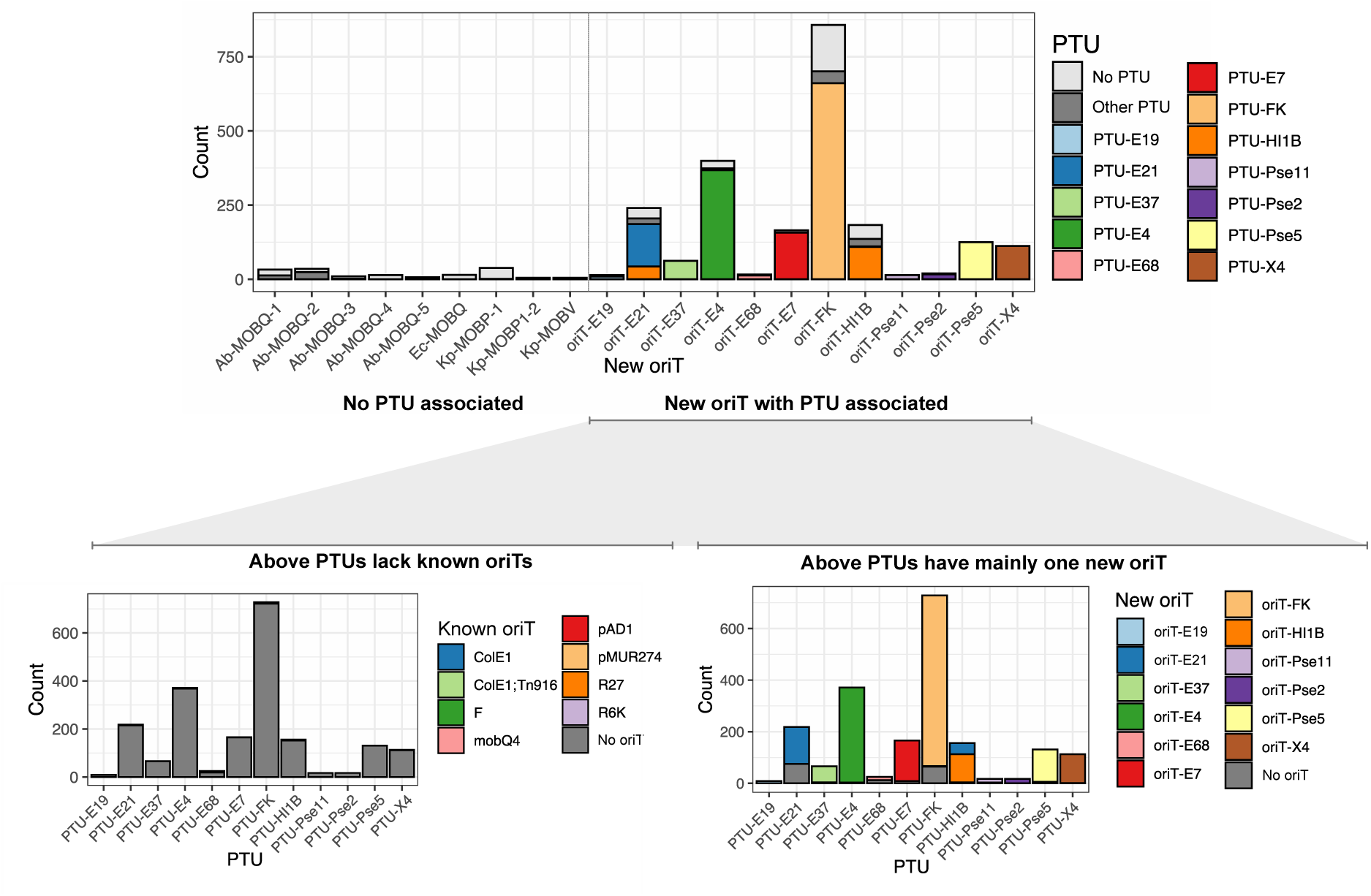
In the upper plot, the Plasmid Taxonomic Units (PTUs) in which the new families of putative *oriTs* were identified. The colors indicate the PTUs, being the legend at the right of the figure. In the bottom plots, PTUs with an associated novel oriT are screened for the presence of known *oriTs* (left) and novel putative ones (right). The colors represent the *oriTs*, being the legend at the right of each plot.

**Figure S11.**
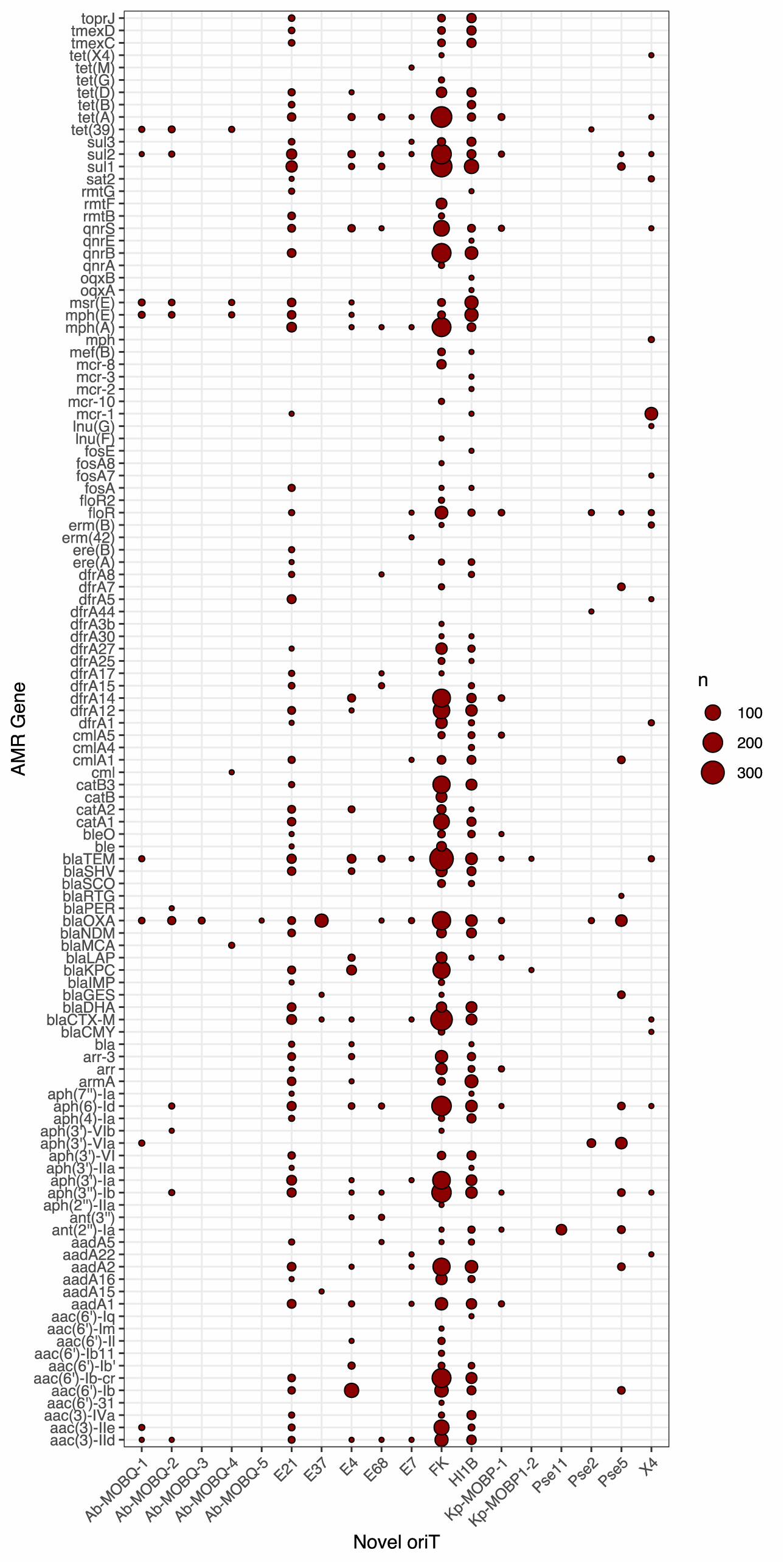
Antimicrobial resistance genes encoded by plasmids with novel families of *oriTs*.

**Figure S12.**
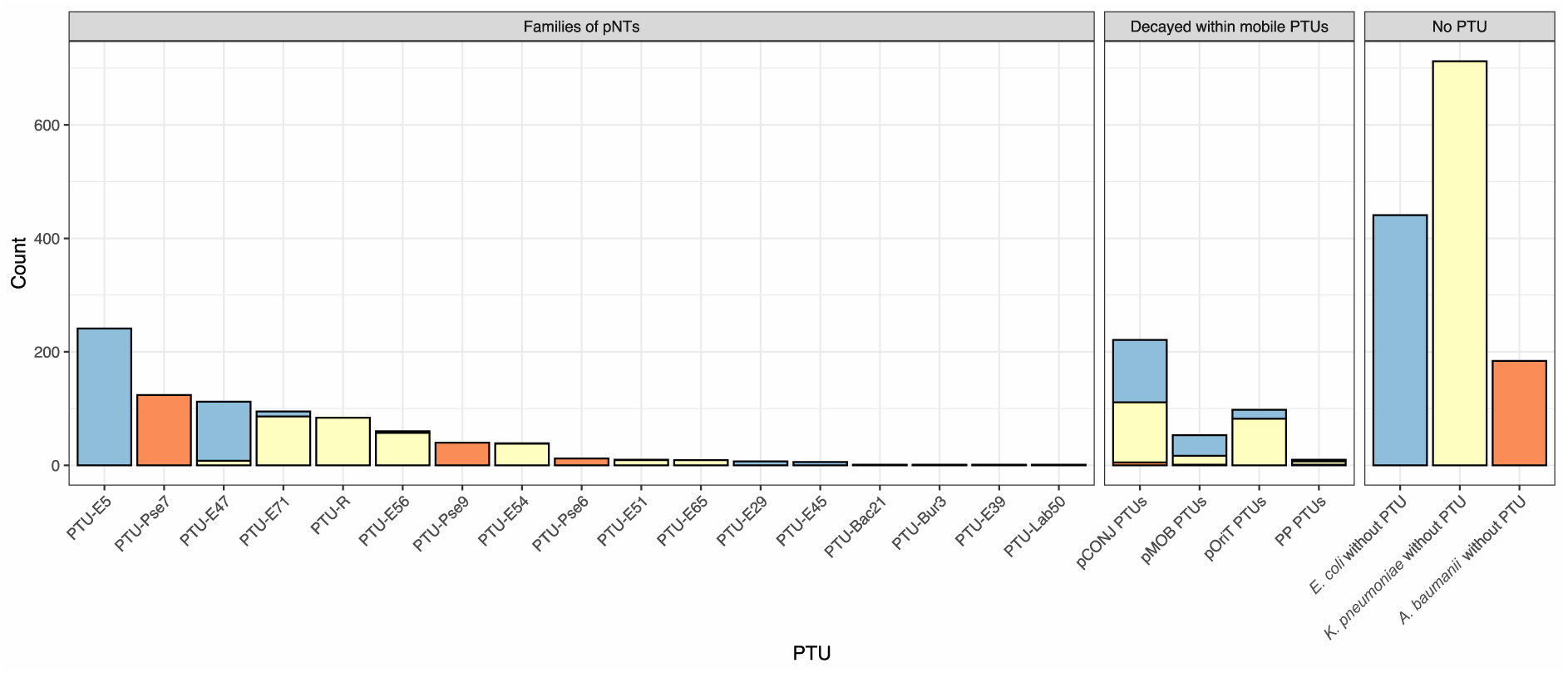
PTUs of the presumably non-transmissible plasmids. “Families of pNTs” represent PTUs formed almost exclusively of pNTs; “Decayed pNTs” represent PTUs in which pNTs are the exceptions; “No PTU”, represent the pNTs with no PTU assigned. The colors represent the species of the plasmid host: *E. coli* (blue), *K. peumoniae* (yellow), *A. baumannii* (orange).

**Figure S13.**
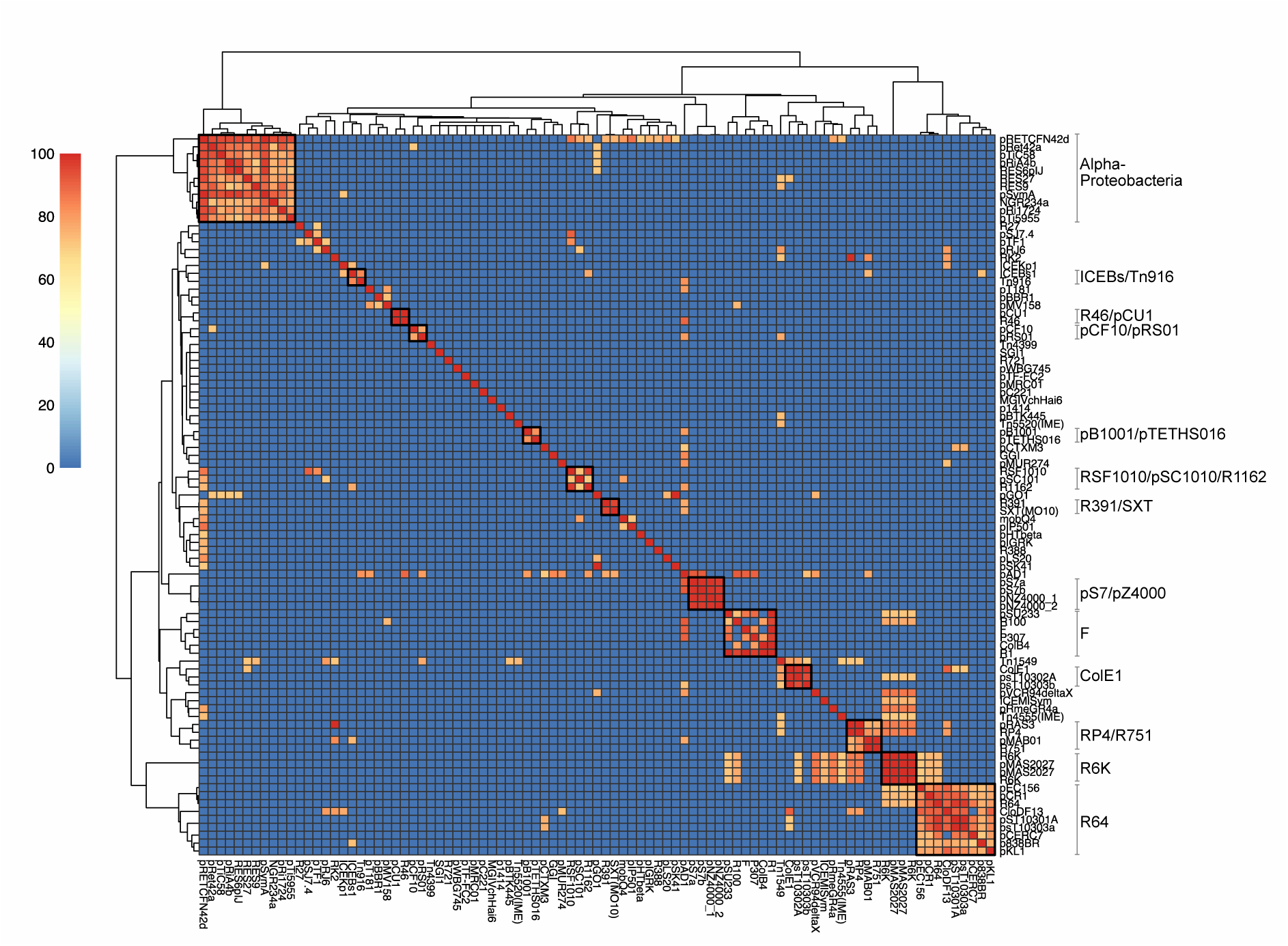
Heatmap of the identity (%) of the all-vs-all local alignments between the origins of transfer of the collection. The black squares within the plot represent the 13 *oriT* families obtained in the analysis. The name of these *oriT* families and the *oriTs* they include is shown at the right of the plot.

**Figure S14.**
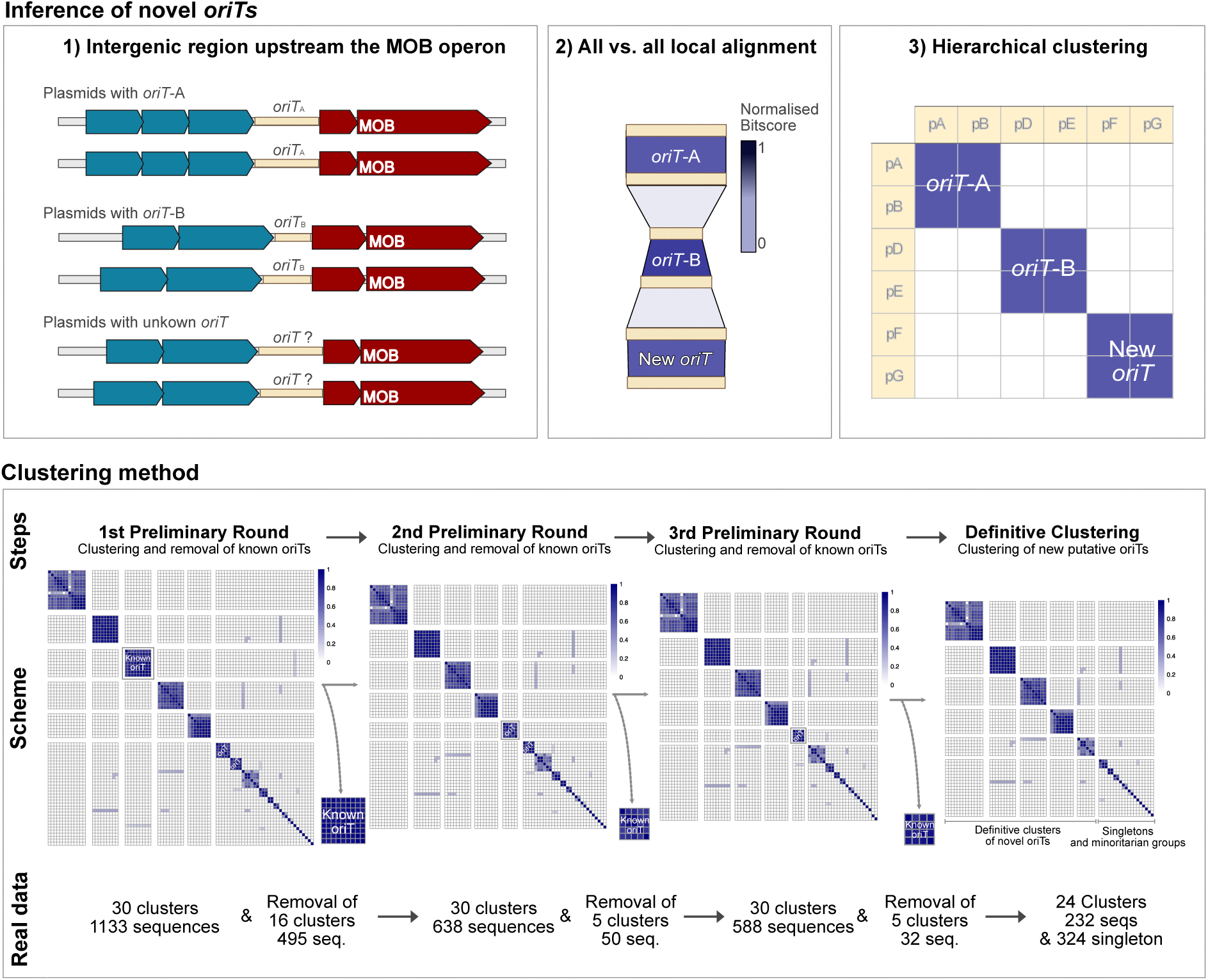
The upper panel represents the method used for the discovery of ovel families of *oriTs*: (1) retrieval of the first non-coding sequence upstream the relaxases. (2) All vs. all local alignments of the non-coding sequences. (3) Clustering of the non-coding sequences using the bitscore of the alignments. The bottom plots represent the clustering method. Three steps of hierarchical clustering prior to the definitive clustering of unknown-*oriT*-carrying sequences. The steps, schematic representation of heatmaps with non-real data, and the real numbers are represented.

